# A force-sensitive mutation reveals a spindle assembly checkpoint-independent role for dynein in anaphase progression

**DOI:** 10.1101/2023.08.03.551815

**Authors:** David Salvador-Garcia, Li Jin, Andrew Hensley, Mert Gölcük, Emmanuel Gallaud, Sami Chaaban, Fillip Port, Alessio Vagnoni, Vicente José Planelles-Herrero, Mark A. McClintock, Emmanuel Derivery, Andrew P. Carter, Régis Giet, Mert Gür, Ahmet Yildiz, Simon L. Bullock

## Abstract

The cytoplasmic dynein-1 (dynein) motor organizes cells by shaping microtubule networks and moving a large variety of cargoes along them. However, dynein’s diverse roles complicate *in vivo* studies of its functions significantly. To address this issue, we have used gene editing to generate a series of missense mutations in *Drosophila* Dynein heavy chain (Dhc). We find that mutations associated with human neurological disease cause a range of defects in larval and adult flies, including impaired cargo trafficking in neurons. We also describe a novel mutation in the microtubule-binding domain (MTBD) of Dhc that, remarkably, causes metaphase arrest of mitotic spindles in the embryo but does not impair other dynein-dependent processes. We demonstrate that the mitotic arrest is independent of dynein’s well-established roles in silencing the spindle assembly checkpoint. *In vitro* reconstitution and optical trapping assays reveal that the mutation only impairs the performance of dynein under load. *In silico* all-atom molecular dynamics simulations show that this effect correlates with increased flexibility of the MTBD, as well as an altered orientation of the stalk domain, with respect to the microtubule. Collectively, our data point to a novel role of dynein in anaphase progression that depends on the motor operating in a specific load regime. More broadly, our work illustrates how cytoskeletal transport processes can be dissected *in vivo* by manipulating mechanical properties of motors.

## INTRODUCTION

Microtubule motors play a key role in organizing the intracellular environment. These molecules power the movement of a wide variety of cellular constituents along microtubules (Hirokawa *et al*, 2009; Reck-Peterson *et al*, 2018) and provide pushing and pulling forces that shape microtubule networks (Cross & McAinsh, 2014; Laan *et al*, 2012; Lu & Gelfand, 2017).

The cytoplasmic dynein-1 (dynein) motor is responsible for almost all microtubule minus-end-directed motor activity in the cytoplasm. Dynein has six subunits – a heavy chain, an intermediate chain, a light intermediate chain and three light chains – which are each present in two copies per complex (Figure 1A). The heavy chain has over 4600 amino acids and comprises an N-terminal tail domain, which mediates homodimerization and association with the other subunits, and a C-terminal motor domain. The key elements of the motor domain are a ring of six AAA+ ATPase domains, a linker, and an anti-parallel coiled-coil stalk that leads to a globular microtubule-binding domain (MTBD) (Figure 1A; Schmidt & Carter, 2016). The major ATP hydrolysis site in dynein is located in the first AAA+ domain (AAA1). ATP binding at this site triggers conformational changes in the ring that are transmitted, via a shift in the stalk’s coiled-coil registry, to the MTBD. This event lowers the affinity of the MTBD for the microtubule, leading to its detachment. ATP hydrolysis and product release allow the MTBD to rebind the microtubule, initiating a swing of the linker that produces force. Repeated cycles of these events drive the movement of dynein along the microtubule. Dynein’s force output and processivity are greatly stimulated by simultaneous binding of two co-factors to the tail domain: another large, multi-subunit complex called dynactin, and a long, coiled-coil-containing cargo adaptor, which is termed an activating adaptor (Belyy *et al*, 2016; McKenney *et al*, 2014; Olenick & Holzbaur, 2019; Reck-Peterson *et al*, 2018; Schlager *et al*, 2014).

**Figure 1.**
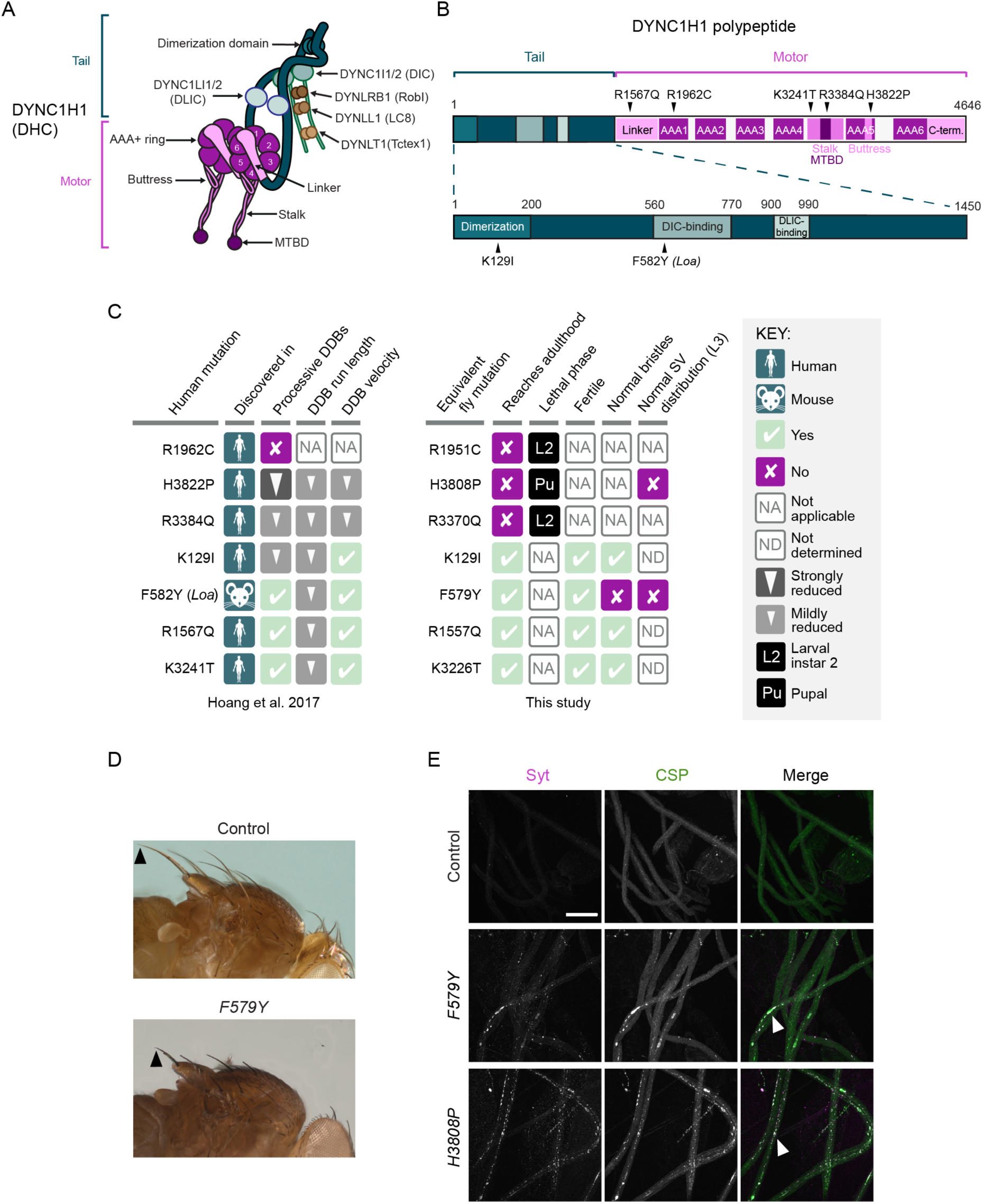
Dynein organization and phenotypic analysis of Dhc disease-associated mutations in *Drosophila*. (A) Cartoon of human dynein complex with alternative nomenclature for subunits shown. MTBD, microtubule-binding domain. The C-terminal domain of DYNC1H1 is not visible in this view as it lies on the other face of the AAA+ rings. (B) Positions in the human DYNC1H1 polypeptide of the disease-associated mutations characterized in this study. Subdomains of the DYNC1H1 polypeptide are color-coded as in panel A. The mouse *Loa* mutation is numbered according to the equivalent residue in human DYNC1H1. Adapted from Hoang *et al*, 2017. (C) Summary of *in vitro* and *in vivo* effects of disease-associated mutations. DDB, dynein-dynactin-BICD2N; SV, synaptic vesicle; L3, larval instar 3. *In vitro* effects refer to observations when both copies of DYNC1H1 in the dynein complex contain the mutation; *in vivo* phenotypes refer to the homozygous condition. (D) Images showing short bristles on the notum of homozygous *Dhc^F579Y^* adult flies compared to controls (*yw*). Arrowheads point to posterior scutellar macrochaetae as an example. The bristle phenotype of *Dhc^F579Y^* flies was completely penetrant (>160 flies examined). (E) Confocal images of segmental nerves (taken proximal to the ventral ganglion; anterior to the top; Z-projections) from fixed L3 larvae stained for the synaptic vesicle proteins Synaptotagmin (Syt) and Cysteine-string protein (CSP). Arrowheads show examples of synaptic vesicle accumulations in mutants. Images are representative of 3 – 6 larvae analyzed per genotype. Scale bar: E, 50 µm.

The wide variety of dynein functions in cells makes it very challenging to assess the function of the motor in discrete processes. Mutations that strongly disrupt dynein function are typically not compatible with survival (Gepner *et al*, 1996; Harada *et al*, 1998; Mische *et al*, 2008) and acute inhibition of the motor complex with function-blocking antibodies (e.g. Gaglio *et al*, 1997; Sharp *et al*, 2000; Yang *et al*, 2007; Yi *et al*, 2011) or small molecule inhibitors (e.g. Firestone *et al*, 2012; Hoing *et al*, 2018; Steinman *et al*, 2017) impairs multiple processes that can be difficult to disentangle.

A striking illustration of the complexity of studying dynein function is in mitosis. Here, the motor has been implicated in centrosome separation (Gonczy *et al*, 1999; Raaijmakers *et al*, 2012; Robinson *et al*, 1999; Vaisberg *et al*, 1993; van Heesbeen *et al*, 2014), nuclear envelope breakdown (Salina *et al*, 2002), spindle pole focusing (Borgal & Wakefield, 2018; Gaglio *et al*, 1997; Merdes *et al*, 1996), attachment of kinetochores to microtubules and chromosome congression (Barisic & Maiato, 2015; Gassmann *et al*, 2008; Li *et al*, 2007; Varma *et al*, 2008; Yang *et al*, 2007), creating tension between sister kinetochores (Sivaram *et al*, 2009; Varma *et al*, 2008; Yang *et al*, 2007), and silencing the spindle assembly checkpoint (SAC) by stripping regulatory factors from kinetochores (Gassmann *et al*, 2010; Griffis *et al*, 2007; Hinchcliffe & Vaughan, 2017; Howell *et al*, 2001; Sivaram *et al*, 2009; Wojcik *et al*, 2001). It was proposed that dynein is also directly responsible for segregating chromosomes in anaphase (Savoian *et al*, 2000; Sharp *et al*, 2000; Yang *et al*, 2007). However, other proteins have since been shown to be essential for poleward chromosome movement and it has been suggested that delays in this process when dynein is inhibited arise indirectly from perturbing the motor’s other functions at the kinetochore (Bader & Vaughan, 2010; Maddox *et al*, 2002; Vukusic *et al*, 2019).

In this study, we have taken a combined genetic and biochemical approach to dissect dynein’s diverse functions. We first document organismal and cellular effects of a large number of Dynein heavy chain missense mutations – including several that are associated with human neurological disease – in the fruit fly, *Drosophila melanogaster*. We then focus on a novel mutation in the MTBD – S3372C – that specifically blocks mitosis in the early embryo. We show that this mutation prevents progression of mitotic spindles from metaphase independently of dynein’s well-established function in licensing anaphase onset by silencing the SAC. High-resolution optical trapping of *in vitro* reconstituted dynein-dynactin-activating adaptor complexes reveals that the MTBD mutation only impairs dynein performance when it encounters resistive loads. All-atom molecular dynamics (MD) simulations show that the S3372C change causes dynein to bind to the microtubule with enhanced flexibility and an altered relative orientation of the stalk. Based on these observations, we propose that dynein has a non-canonical function in anaphase progression that involves the motor operating in a specific force regime. Our work illustrates the feasibility of dissecting the function of cytoskeletal motors by interfering with specific mechanical properties.

## RESULTS

### Characterization of disease-associated Dhc mutations in *Drosophila*

We first set out to study the *in vivo* effects of six neurological-disease-linked missense mutations in the human Dynein heavy chain protein (DYNC1H1): K129I, R1567Q, R1962C, R3384Q, K3241T and H3822P (Figure 1B). Heterozygosity for each of these mutations is associated with malformations in cortical development and intellectual disability (Poirier *et al*, 2013; Schiavo *et al*, 2013; Vissers *et al*, 2010; Willemsen *et al*, 2012). We previously used *in vitro* reconstitution to show that these mutations perturb, to varying degrees, the processive movement of human dynein complexes bound to dynactin and the N-terminal region of the prototypical activating adaptor, BICD2 (so-called ‘DDB’ complexes) (Hoang *et al*, 2017; results summarized in Figure 1C). Whether these inhibitory effects accounted fully for the *in vivo* consequences of these mutations was unclear.

We therefore used CRISPR/Cas9-based homology-directed repair (HDR) to generate *Drosophila* strains with equivalent mutations in the gene encoding Dynein heavy chain (*Dhc64C*, hereafter *Dhc*) (Figure 1C). We also made a strain carrying the *Drosophila* equivalent of the human F582Y mutation (F579Y), which corresponds to the *Legs-at-odd-angles* (*Loa*) allele that causes neurodegenerative disease in heterozygous mice (Hafezparast *et al*, 2003). This mutation also impairs the movement of DDB complexes along microtubules *in vitro* (Hoang *et al*, 2017; Figure 1C).

None of the mutations caused overt phenotypes when heterozygous in flies. However, animals homozygous for three of the mutations (R1962C human/R1951C fly; H3822P human/H3808P fly; R3384Q human/R3370Q fly) failed to reach adulthood (Figure 1C). Complementation tests with a *Dhc* protein null allele (Fumagalli *et al*, 2021) confirmed that lethality was due to the missense mutations rather than off-target activity of CRISPR/Cas9 (Figure S1A). Whereas homozygous *Dhc* protein null mutants died during both early and late phases of the second larval instar stage (L2), the vast majority of *R1951C* and *R3370Q* mutants arrested during early L2 (Figure 1C and Figure S1B). The slightly earlier lethal phase of R1951C and R3370Q compared to the *Dhc* null suggests that these mutations have a partially dominant negative effect. *H3808P* mutants typically died during the pupal phase (Figure 1C and Figure S1B), demonstrating that this mutation retains some dynein activity *in vivo*. Of the four homozygous viable mutations (K129I, F579Y, R1557Q (equivalent to R1567Q) and K3226T (equivalent to human K3241T)), only F579Y caused a morphological defect in adults. Homozygotes for this mutation had abnormally short bristles on the notum (Figure 1D), which is a feature of several classical hypomorphic *Dhc* mutations (Gepner *et al*, 1996; Melkov *et al*, 2016).

We previously proposed that the pathomechanism of the disease-associated mutations involves impaired trafficking of cargo-motor complexes in neurons (Hoang *et al*, 2017). To investigate this notion, we examined the effects of the H3808P and F579Y mutations on the distribution of synaptic vesicles in axons of segmental nerve motor neurons in larval instar 3 (L3). Synaptic vesicles in this system undergo dynein-and kinesin-1-dependent bidirectional transport (Miller *et al*, 2005), and impairing the activity of either motor causes focal accumulations of the vesicles that are visible by immunofluorescence (Martin *et al*, 1999). Homozygosity for the *H3808P* or *F579Y* alleles caused striking synaptic vesicle accumulations in axons (Figure 1E), supporting the hypothesis that disease-associated mutations interfere with cargo translocation in neurons.

Figure 1C summarizes the effects of the disease-associated mutations in *Drosophila* and on DDB activity *in vitro* (Hoang *et al*, 2017). The three mutations that had the strongest effects on DDB behavior *in vitro* – R1962C, R3384Q, and H3822P – were those whose counterparts caused lethality in flies. However, the relative strengths of these mutations’ *in vitro* and *in vivo* effects were not equivalent. Even though H3822P impaired *in vitro* motility of DDB more strongly than R3384Q, the equivalent mutation resulted in later lethality in flies than the R3384Q equivalent. The milder phenotype of the H-to-P mutation in *Drosophila* was particularly surprising as this change strongly disrupts dynein function in budding yeast (Marzo *et al*, 2019). Moreover, whereas the effect of the human F582Y mutation on DDB *in vitro* motility was very similar to that of R1567Q and K3241T, and weaker than that of K129I, the equivalent *Drosophila* mutation (F579Y) was unique amongst this group in disrupting bristle morphology. These observations might reflect some mutations having species specific effects or altering dynein’s interplay with proteins in addition to dynactin and the BICD2 orthologue, BicD. Nonetheless, the range of mutant phenotypes observed in *Drosophila* and the overall correlation with the strength of inhibitory effects *in vitro* indicate that the fly is a valuable model for structure-function studies of the dynein complex.

### The novel mutation S3372C specifically impairs a function of dynein during embryonic development

Whilst generating the disease-associated mutations in *Drosophila* we recovered four novel in-frame Dhc mutations that were caused by imprecise repair of CRISPR/Cas9-induced double strand breaks. L1952K and the two-amino-acid deletion ΔL3810Y3811, which came from the experiments that produced R1951C and H3808P, respectively, caused lethality in homozygotes (Figure S2A) or when combined with the null allele (Figure S1A). Homozygosity for each of these mutations resulted in arrest during L3 and accumulations of synaptic vesicles in motor neuron axons (Figure S2A and B). Y3811F was also recovered whilst making H3808P, although the homozygous mutants did not have a detectable phenotype (Figures S1A and S2A).

The fourth novel mutation, S3372C, was a by-product of our efforts to make R3370Q. This mutation, which is situated in the MTBD, had a particularly striking phenotype. Whilst *S3372C* homozygous adults were recovered at the normal Mendelian frequency and males were fertile, homozygous females had very strongly impaired fertility (0.029% of eggs hatched into larvae; Figure 2A). No fertility problems were observed in females heterozygous for the *S3372C* allele, showing that this is a recessive effect. Like *S3372C* homozygotes, *trans*-heterozygotes for the *S3372C* allele and the *Dhc* protein null allele (*S3372C/-*) had normal survival to adulthood (Figure S1A) and female-specific infertility (Figure 2A). Thus, the female fertility defect is associated with the S3372C mutation rather than an off-target effect of CRISPR/Cas9. This conclusion was corroborated by the ability of a wild-type *Dhc* genomic construct to restore fertility to *S3372C* homozygotes (Figure S2C).

**Figure 2.**
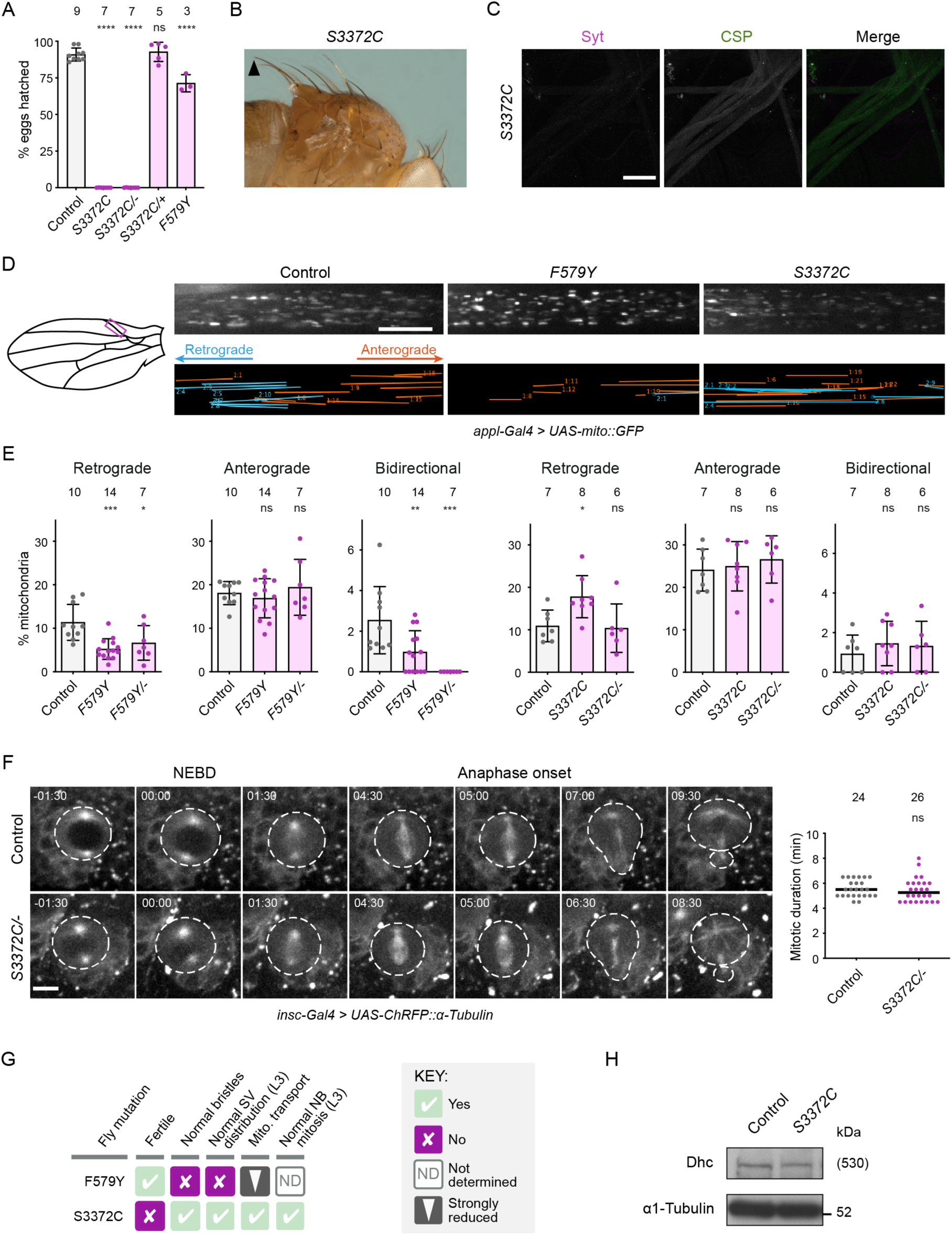
The novel mutation S3372C selectively affects embryonic development. (A) Hatching rate of eggs laid by mated females of the indicated genotypes. Columns show mean values per egg collection; error bars represent S.D.; circles are values for individual egg collections. Number of collections per genotype (from 3 independent crosses) is shown above columns (114 – 575 eggs per collection). *S3372C/-* are *trans*-heterozygous for *S3372C* and a *Dhc* null allele. Control genotype: *yw*. (B, C) Images showing (B) normal bristle length in adults and (C) lack of synaptic vesicle accumulations in segmental nerves of L3 larvae (proximal to the ventral ganglion; anterior to the top; Z-projection) in *S3372C* homozygotes (for comparisons with controls, see Figure 1D and E). Arrowhead in B points to posterior scutellar macrochaetae. Images in B and C are representative of >160 flies and 3 larvae analyzed, respectively. (D) Analysis of mitochondrial motility in wing nerve axons of 2-day-old wild-type, *F579Y* and *S3372C* adult flies. Left, cartoon of wing with magenta box indicating region imaged. Top images, example stills from 3-minute time series (single focal plane) of fluorescent mitochondria (expression of mito::GFP in neurons using *Appl-Gal4*). Bottom images, example traces of motile mitochondria in corresponding time series. Blue, retrograde tracks; orange, anterograde tracks. (E) Percentages of mitochondria transported in the retrograde or anterograde directions, or moving bidirectionally during the 3 minutes of data acquisition. Columns show mean values per movie; error bars represent S.D.; circles are values for individual movies (each from a different wing). Number of wings analyzed shown above bars. Note that we observed an increased frequency of retrograde transport in *S3372C* homozygotes but this was not recapitulated in *S3372C/-* animals. (F) Analysis of mitotic duration in neuroblasts (NBs) in the brain of control and *S3372C/-* mutant L3 larvae. Left, stills from image series of NBs with fluorescently-labeled spindles (expression of RFP-tagged α-tubulin in NBs using *insc-Gal4*). NBs and daughter cells highlighted with dashed line. Timestamps are min:s after nuclear envelope breakdown (NEBD). Right, quantification of mitotic duration (NEBD to anaphase onset). Lines show median; circles are values for individual NBs. Numbers of neuroblasts analyzed (from 4 wild-type or 5 *S3372C/-* larvae) is shown above plot. (G) Summary of *in vivo* effects of homozygosity for *F579Y* and *S3372C*. SV, synaptic vesicle; Mito., mitochondrial. (H) Immunoblot of extracts of embryos from control and *S3372C* mothers (0 – 160 min collections), probed with antibodies to Dhc and α1-tubulin (loading control). Position of molecular weight (Mw) marker is shown for α1-tubulin blot segment; as there is no marker where Dhc migrates, the predicted Mw of Dhc is shown in parentheses. Evaluation of statistical significance (compared to control) was performed with a 1-way ANOVA with Dunnett’s multiple comparisons tests (A and E) or a Mann-Whitney test (F): ****, *P*<0.0001; ***, P<0.001; **, P<0.01; *, P<0.05; ns, not significant. Scale bars: C, 50 µm; D, 10 µm; F, 5 µm.

In contrast to S3372C, the non-lethal disease-associated *Dhc* mutations did not strongly affect fertility (Figure 1C). For example, only 30% of embryos from *F579Y* homozygous mothers failed to hatch (Figure 2A). Moreover, despite having much stronger fertility defects, *S3372C* homozygous and *S3372C/-* animals did not have the bristle defects (Figure 2B and Figure S2D) or focal accumulations of synaptic vesicles (Figure 2C and Figure S2E) observed with the F579Y mutation. These observations reveal that S3372C only affects a subset of dynein functions.

To further explore which dynein-dependent processes are affected by S3372C, we used time-lapse imaging to compare trafficking of GFP-labeled mitochondria in wing nerve axons of adult wild-type and mutant flies (Vagnoni *et al*, 2016). In these axons, as in others, dynein and kinesin-1 are responsible for retrograde and anterograde mitochondrial movements, respectively (Vagnoni *et al*, 2016). The F579Y mutation was used as a positive control for the experiments, as the equivalent mutation impairs dynein-based cargo transport in other organisms (Hafezparast *et al*, 2003; Ori-McKenney *et al*, 2010; Sivagurunathan *et al*, 2012). Homozygosity for *F579Y* allele reduced the frequency of retrograde and bidirectional transport events of mitochondria in wing nerve axons without impairing retrograde velocity or travel distance, or any aspect of anterograde motion (Figure 2D and E and Figure S2F and G). In contrast, the S3372C mutation did not significantly impair any aspect of retrograde or anterograde motion of mitochondria (Figure 2D and E and Figure S2F and G). We also assessed the effect of S3372C on cell division of neuroblasts of live L3 brains expressing fluorescent α-tubulin. Inhibiting dynein and dynactin function in these neuroblasts perturbs spindle morphology and delays the completion of mitosis (Siller *et al*, 2005; Wojcik *et al*, 2001). In *S3372C/-* mutant neuroblasts, however, no spindle abnormalities were observed and the duration of mitosis was indistinguishable from the wild type (Figure 2F). Thus, the S3372C mutation has no discernible inhibitory effect on dynein functions in axonal transport of mitochondria or mitosis of larval neuroblasts.

Taken together, our phenotypic analyses (summarized in Figure 2G) reveal a selective maternal effect of S3372C on dynein function during embryogenesis. A trivial reason for this observation could be that the mutation destabilizes Dhc protein in the embryo. However, immunoblotting of wild-type and *S3372C* embryo extracts with an α-Dhc antibody showed that this is not the case (Figure 2H). Therefore, the MTBD mutation affects the functionality, rather than the level, of Dhc in embryos.

### The conserved location of S3372 indicates an important role in MTBD function

To attempt to gain insight into how mutating S3372 affects dynein activity, we examined its position within the tertiary structure of the MTBD. Whilst an experimentally determined structure of the *Drosophila* dynein MTBD is not available, we could confidently infer the position of the serine from a 4.1 Å-resolution cryo-electron microscopy (EM) structure of the closely related mouse MTBD, together with a portion of the stalk, bound to an α/β-tubulin dimer in the microtubule (Figure 3A; Lacey *et al*, 2019). The equivalent residue in mouse (S3384) is located in a short loop that connects helix 6 (H6) of the MTBD – which directly contacts α-tubulin – to the base of coiled-coil 2 (CC2) of the stalk (Figure 3A and B). We also used Alphafold2 (Jumper *et al*, 2021) to produce high-confidence structural predictions of the MTBD and stalk of the *Drosophila* and human dynein-1 heavy chains (Figure 3B and Figure S3A). In both these structures, the serine adopted an equivalent position to that observed in the cryo-EM structure of the mouse MTBD.

**Figure 3.**
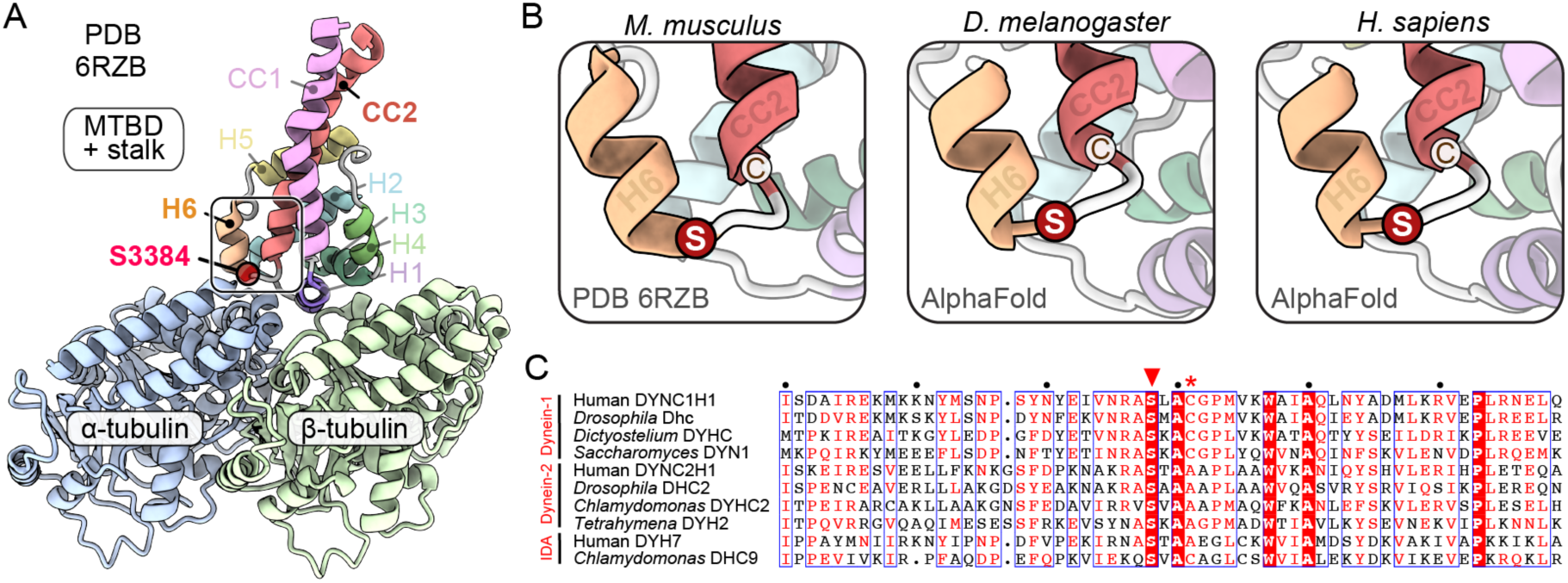
S3372 has a conserved position in a loop between H6 of the MTBD and CC2 of the stalk. (A) Overview of position of the equivalent residue to S3372 (S3384) in the cryo-EM structure of the mouse MTBD and portion of the stalk in complex with the α/β-tubulin dimer (PDB, 6RZB). Position of S3384 is highlighted by a red circle. (B) Zoom ins of regions containing S3384 in 6RZB and equivalent residues (red circles) in Alphafold2-generated structures of the MTBD and stalk of *Drosophila melanogaster* Dhc and human DYNC1H1. Neighbouring cysteine residues (C3387 mouse; C3375 *Drosophila*; C3389 human) are also shown. (C) Alignment of sequences from the MTBD and CC2 of the stalk of the indicated dynein family members. White letters on a red background, residues present in all sequences; red letters, residues present in ≥ 50% of sequences; blue boxes, regions with ≥ 50% conservation; red arrowhead, residues equivalent to S3372 of *Drosophila* Dhc; red asterisk, residues equivalent to C3375 of *Drosophila* Dhc. Uniprot accession numbers: human (*Homo sapiens*) DYNC1H1, Q14204; *Drosophila melanogaster* Dhc, P37276; *Dictyostelium discoideum* DYHC, P34036; *Saccharomyces cerevisiae* DYN1, P36022; human (*Homo sapiens*) DYNC2H1, Q8NCM8; *Drosophila melanogaster* DHC2, Q0E8P6; *Chlamydomonas reinhartii* DYHC2, Q9SMH5; *Tetrahymena thermophila* DYH2, Q5U9X1; human (*Homo sapiens*) inner dynein arm (IDA) DYH7, Q8WXX0; *Chlamydomonas reinhartii* inner dynein arm (IDA) DHC9, Q4AC22.

The conserved position of the serine in dynein-1 from fly, mouse and human suggests that it plays an important role in these contexts. To determine if this residue could have broader significance, we examined its conservation in other dynein-1 proteins, as well as in axonemal dyneins and dynein-2, which drive the beating of cilia and movement of material within these structures, respectively (Antony *et al*, 2021). The serine was conserved in the primary sequence of all dynein-1, dynein-2, and axonemal dynein heavy chain sequences examined, including those from single cell eukaryotes (Figure 3C). Alphafold2-generated predictions of MTBD and stalk regions from the human dynein-2 heavy chain and axonemal DYH7 proteins (Figure S3A) indicated that the location of the serine in a loop between H6 and CC2 is also conserved (Figure S3B). These analyses suggest that the serine that is mutated in the *S3372C* fly strain plays an important role across the dynein family.

### Ectopic disulfide bond formation does not account for the S3372C phenotype

The above structural analysis also revealed that, whilst there is no nearby cysteine on the α/β-tubulin dimer, there is a conserved cysteine at the base of CC2 in dynein-1 proteins (C3375 in *Drosophila*) that is in close proximity to the serine mutated in the *S3372C* strain (Figure 3B and C). These observations led us to hypothesize that the S3372C mutation results in an ectopic disulfide bond between this residue and C3375. There is a global increase in oxidation at the *Drosophila* egg-to-embryo transition (Petrova *et al*, 2018), which could conceivably trigger disulfide bonding at this site. Such an event could impair dynein’s dynamics specifically in the embryo and thereby explain arrest at this stage. To test this notion, we used CRISPR/Cas9-mediated HDR in *Drosophila* to change S3372 to cysteine and C3375 to serine in the same polypeptide, thus eliminating the potential for a disulfide bond at these positions (Figure 4A). Like *S3372C* females, females homozygous for this allele (*S3372C + C3375S*) were almost completely infertile (Figure 4B). This effect was not associated with the C3375S change, as females with this mutation had only modestly reduced fertility (Figure 4B). Thus, disulfide bonding with C3375 does not account for the embryonic arrest caused by S3372C.

**Figure 4.**
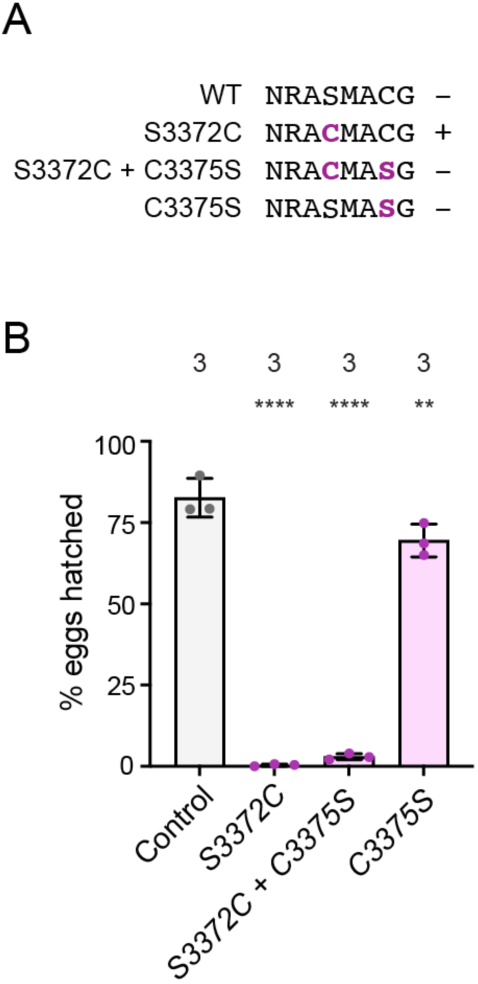
The S3372C embryonic arrest phenotype is not caused by ectopic disulfide bonding in Dhc. (A) Sequences of relevant regions of wild-type (WT), S3372C, S3372C + C3375S, and C3375S *Drosophila* Dhc. Mutated residues are shown in magenta. Potential for intramolecular disulfide bond formation is indicated with a plus. (B) Quantification of hatching rate of eggs laid by mated females of the indicated genotypes. Columns show mean values per egg collection; error bars represent S.D.; circles are values for individual egg collections. Number of collections per genotype (each from an independent cross; 245 – 736 eggs per collection) is shown above bars. Control genotype was homozygous for a wild-type *Dhc* allele recovered from the same CRISPR-Cas9 mutagenesis experiment that generated the *Dhc* mutant alleles. Evaluation of statistical significance (compared to control) was performed with a 1-way ANOVA with Dunnett’s multiple comparisons test: ****, P<0.0001; **, P<0.01.

### S3372C causes metaphase arrest of early nuclear divisions

Our finding that S3372C does not destabilize Dhc protein in the embryo or cause ectopic disulfide bonding points to a more nuanced effect on dynein activity, possibly related to a specific function during development. To elucidate which motor function is affected by this mutation, we investigated the embryonic phenotype in more detail. DNA staining of fixed embryos revealed that the vast majority of embryos from *S3372C* homozygous mothers did not reach syncytial blastoderm stages, with ∼80% arresting during early nuclear cleavage cycles (Figure 5A and B). Development of *S3372C* embryos to this stage was dependent on fertilization because only a single nucleus was observed in unfertilized mutant eggs (Figure S4A), as is the case in unfertilized wild-type eggs (Loppin *et al*, 2015). Arrest during early cleavage stages was also typical for embryos from *S3372C/-* mothers (Figure S4A), demonstrating the causal nature of the MTBD mutation. In contrast, embryos from *F579Y* mothers developed normally to the blastoderm stage (Figure S4A). The partially penetrant *F579Y* hatching defect observed previously (Figure 2A) must therefore be due to problems arising later in development.

**Figure 5.**
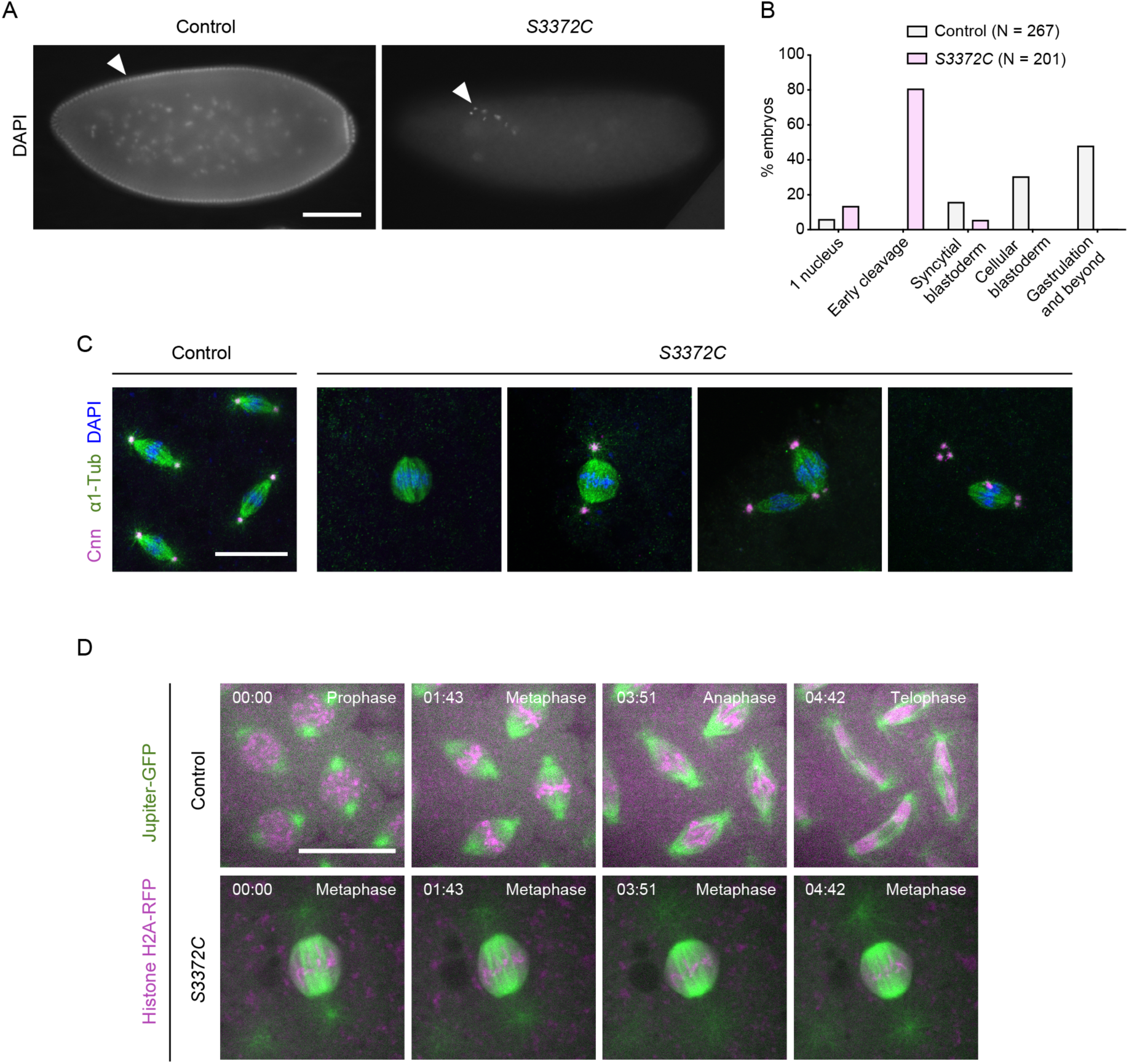
S3372C causes metaphase arrest in the embryo. (A) Example wide-field images of fixed embryos from a 2 to 4-h egg collection from mated control (*yw* strain) and *S3372C* mothers stained with DAPI (arrowheads show DNA staining). (B) Quantification of stages of control and *S3372C* embryos from 2 to 4-h egg collections. N is number of embryos scored. The ‘1 nucleus’ category includes potentially unfertilized eggs. (C) Example confocal images of mitotic spindles in fixed control and *S3372C* embryos (Z-projection). Cnn, Centrosomin (centrosome marker); α1-Tub, α1-Tubulin (microtubules). (D) Example stills from time series (single focal plane) of control and *S3372C* embryos acquired during preblastoderm cycles. Jupiter-GFP and His2Av-mRFP label microtubules and chromatin, respectively. Timestamps are min:s. Scale bars: A, 100 µm; C and D, 20 µm.

To visualize the mitotic apparatus of *S3372C* mutant embryos, we additionally stained them for markers of centrosomes and microtubules. This revealed that the vast majority of mitotic spindles in S3372C embryos were arrested in metaphase (Figure 5C and Figure S4B). Moreover, we found that, whilst 70% of spindles in the mutants had centrosomes, only 10% had the canonical arrangement of one centrosome at each pole (Figure S4B). The other mutant spindles had a range of abnormalities in centrosome arrangement, including missing or supernumerary centrosomes at one pole, or detachment of centrosomes from the spindle (Figure 5C and S4B). Time-lapse analysis of mitosis in embryos that had fluorescently-labeled microtubules in combination with either fluorescently-labeled histones (Figure 5D and Movie S1) or centrosomes (Figure S4C and Movie S2) confirmed that S3372C causes metaphase arrest and defects in centrosome arrangement. The observation that some metaphase-arrested spindles had normal positioning and numbers of centrosomes (Figure S4B) suggested that altered centrosome arrangement is not primarily responsible for the metaphase arrest of *S3372C* spindles. Instead, a failure to progress to anaphase may uncouple mitotic cycles from centrosome duplication cycles, a scenario previously observed upon inhibition of several mitotic regulators (Archambault & Pinson, 2010; Defachelles *et al*, 2015; McCleland & O’Farrell, 2008).

The above analysis shows that the S3372C mutation blocks anaphase onset of mitotic spindles in the early embryo. To investigate if all dynein functions are affected at this stage of development, we examined the maintenance of the mitotic phase (M-phase) of polar bodies. Dynein has been implicated in this process through analysis of embryos in which the function of BicD or another dynein-associated protein, Rod, is disrupted (Defachelles *et al*, 2015; Vazquez-Pianzola *et al*, 2022). In a sizeable fraction of BicD-or Rod-deficient embryos, the polar body is not arrested in metaphase as judged by decondensation of the DNA, as well as the absence of phosphorylated histone H3 and the SAC component BubR1. We saw no such defects in *S3372C* embryos (Figure S5A and B), indicating that dynein’s function in maintaining M-phase of the polar body is not impaired. Thus, the S3372C mutation appears to only affect a subset of dynein functions in the early embryo that include – and are perhaps limited to – the progression of mitotic spindles from metaphase to anaphase.

### S3372C increases dynein accumulation in the vicinity of the kinetochore

To attempt to narrow down how the S3372C mutation causes metaphase arrest of mitotic spindles, we investigated its effect on the distribution of the motor complex on the mitotic apparatus. This was achieved by comparing the localization of a GFP-tagged Dynein light intermediate chain subunit (GFP-Dlic) in embryos laid by wild-type and *S3372C* homozygous females. In wild-type embryos, GFP-Dlic was associated with the spindle for the duration of the mitotic cycle with a transient, weak enrichment at kinetochores during prometaphase and metaphase (Figure 6A and Movie S3). This pattern is in keeping with previous observations of dynactin localization during embryonic mitosis (Wojcik *et al*, 2001). In contrast, all *S3372C* mutant spindles had a very strong accumulation of GFP-Dlic signal in the vicinity of the kinetochores, which often extended along the neighbouring microtubules (Figure 6A and Movie S3). The mutant embryos did not, however, have an overt enrichment of GFP-Dlic on other regions of the spindle apparatus, including the spindle poles or centrosomes (Figure 6A and Movie S3). To test if the abnormal accumulation of dynein at the kinetochore region of mutant embryos is solely an indirect effect of the metaphase arrest, we examined GFP-Dlic localization in embryos of *S3372C/+* mothers, which have no apparent defects in mitosis. Unlike the wild type, a fraction of these embryos (43%) had ectopic kinetochore association of dynein (Figure 6B), albeit less strong than in embryos from *S3372C* homozygous mothers and only at a subset of kinetochores. Collectively, these observations suggest that the S3372C mutation can directly influence kinetochore-related functions of dynein.

**Figure 6.**
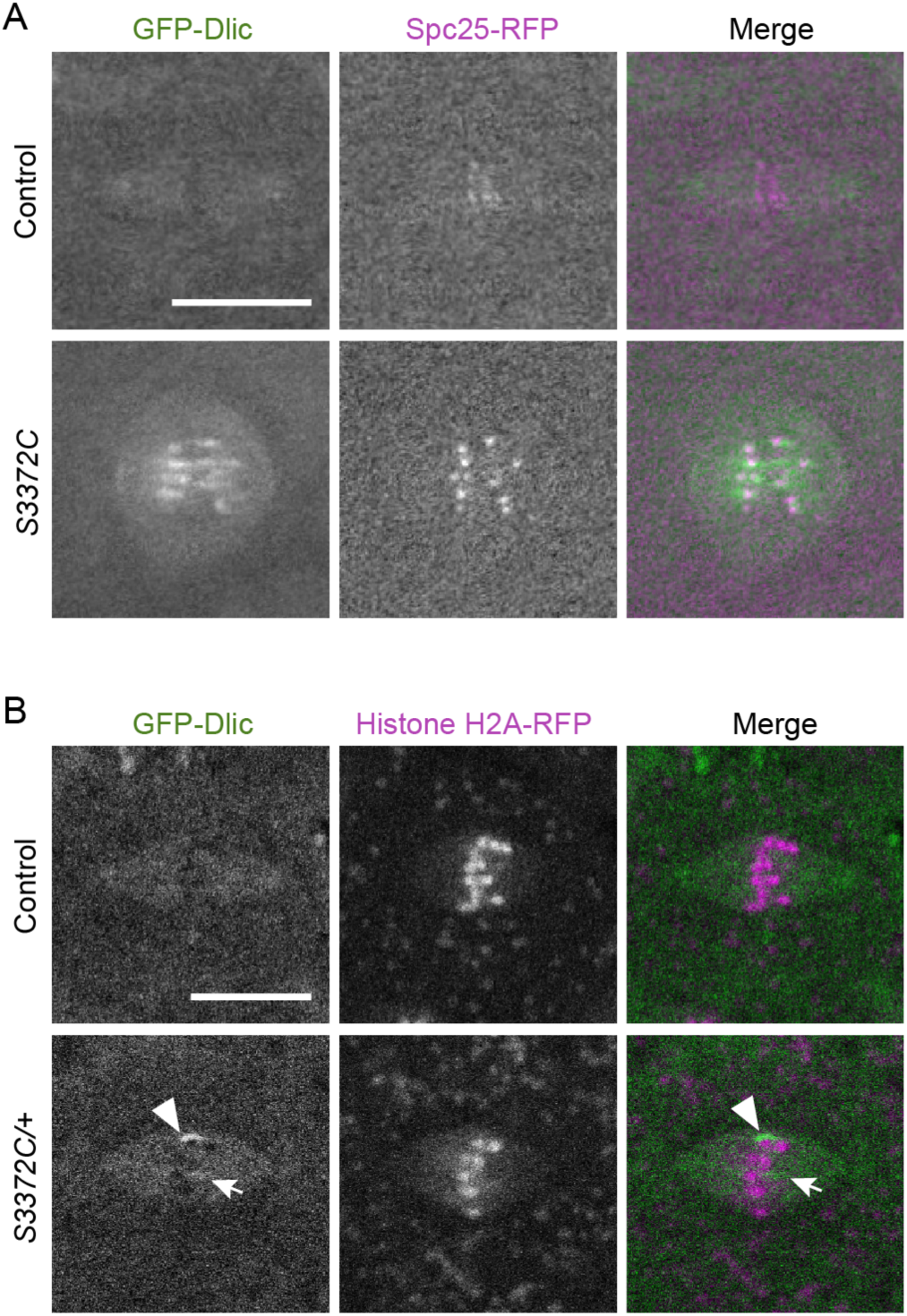
S3372C cause ectopic accumulation of dynein in the vicinity of kinetochores. (A, B) Example confocal images (Z-projections) of mitotic spindles in live control and (A) *S3372C* and (B) *S3372C/+* embryos with fluorescently-labeled dynein (GFP-Dlic). Embryos additionally contain markers of kinetochores (Spc25-RFP; A) and chromatin (Histone H2A-RFP; B). In B, arrow and arrowhead show examples of accumulation of dynein at the metaphase plate in *S3372C/+* embryos. Ectopic GFP-Dlic accumulation at ≥ 1 kinetochore was observed in 100% (21/21) of *S3372C* embryos, 43% (10/23) of *S3372C/+* embryos and 0% (20/20) of control embryos. Scale bars: 10 µm.

### S3372C blocks anaphase progression in a SAC-independent manner

The above findings prompted us to examine kinetochore-associated roles of dynein in the mutant embryos in more detail. Functions of dynein at the kinetochore are closely linked with silencing of the SAC (Gassmann, 2023). This checkpoint delays metaphase-anaphase transition by inhibiting co-activators of the anaphase promoting complex/cyclosome (APC/C) ubiquitin ligase until chromosomes have been successfully biorientated (Musacchio, 2015).

One function of dynein in this context is to generate end-on attachments of kinetochores with microtubule plus ends, which is a prerequisite for SAC silencing (Bader & Vaughan, 2010; Barisic & Maiato, 2015). In *S3372C* embryos, the ends of microtubule bundles were closely apposed with the kinetochore-associated proteins Spc25-RFP (Figure 7A) and the dynein activating adaptor Spindly (Figure S5C). This observation indicates that metaphase arrest in the mutants is not due to a failure of kinetochores to engage with microtubule plus ends. However, with the resolution available within the *Drosophila* embryo it was not possible to tell if the number or orientation of kinetochore-microtubule attachments were established correctly.

**Figure 7.**
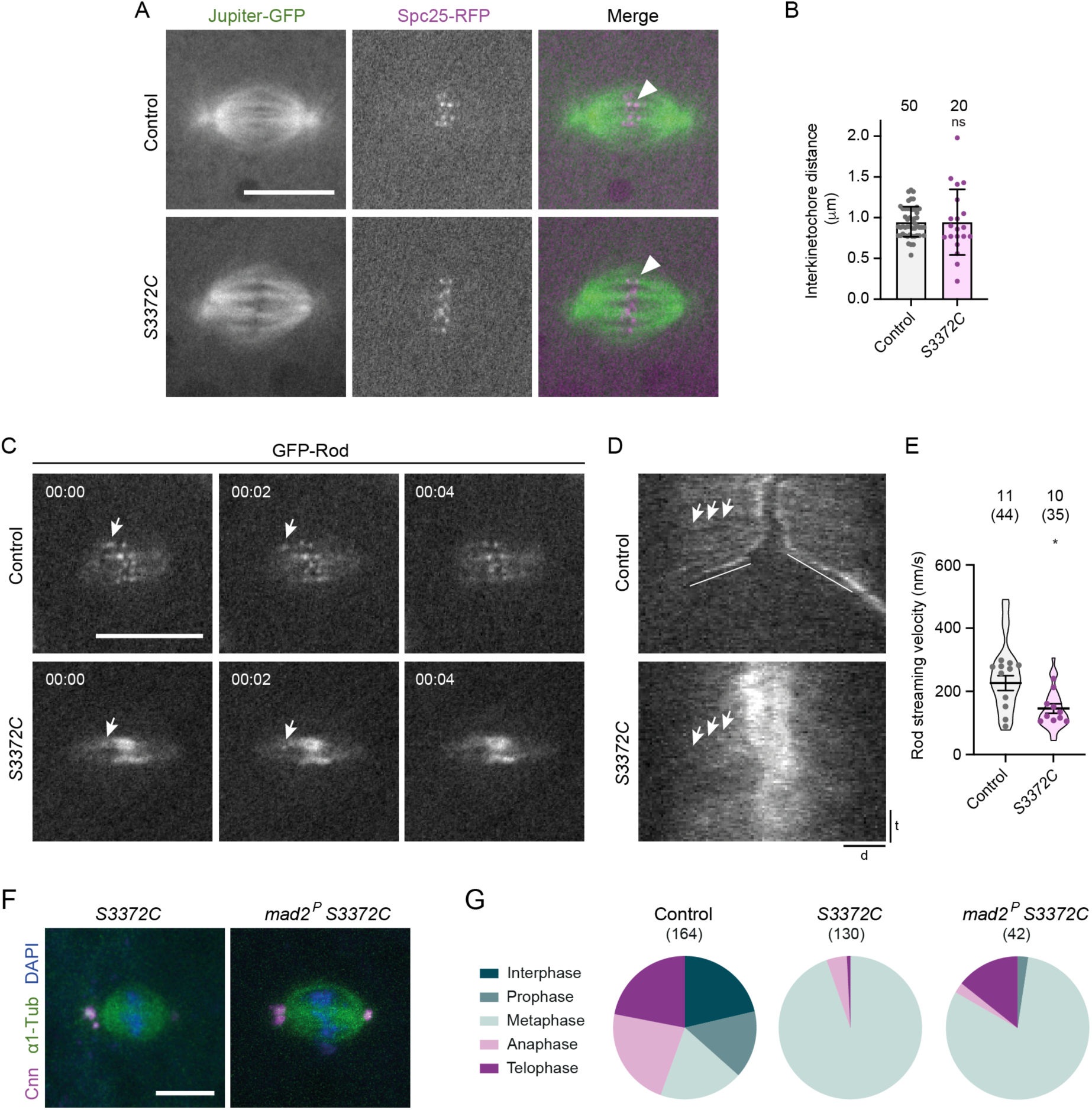
S3372C blocks mitosis in a SAC-independent manner. (A) Example confocal images of mitotic spindles (single focal plane) in live control and *S3372C* embryos that have fluorescently-labeled microtubules (Jupiter-GFP) and kinetochores (Spc25-RFP). Arrowheads, examples of association of bundled microtubules with a kinetochore. (B) Quantification of interkinetochore distance (using centroids of Spc25-RFP signals) in live control and *S3372C* metaphase spindles. Columns show mean values; error bars represent S.D.; circles are values for individual kinetochore pairs. Number of kinetochore pairs analyzed (from 7 control and 8 *S3372C* embryos) is shown above bars. (C) Example stills from a time series (single focal plane) showing streaming (arrows) of GFP-Rod particles in control and *S3372C* embryos. Timestamps are min:s. (D) Example kymographs showing GFP-Rod streaming (e.g. arrows). d, distance; t, time. Diagonal lines in control label movements of kinetochores in anaphase. (E) Quantification of GFP-Rod streaming velocity. Violin plots show values for individual motile Rod particles and circles show mean values per embryo. Horizontal lines shown mean ± S.D. of values for individual motile particles. Numbers without parentheses above bars are number of embryos, with total numbers of particles given in parentheses. (F) Example confocal images of metaphase spindles from fixed embryos of *S3372C* and *mad2^P^, S3372C* mothers stained as indicated (Z-projections). (G) Incidence of mitotic stages in control, *S3372C* and *mad2^P^, S3372C* embryos (25 – 130 minute collection). Numbers in parentheses indicate number of nuclei or spindles counted per genotype; data are from 34 control, 49 *S3372C* and 33 *mad2^P^, S3372C* embryos, with no more than 5 randomly selected nuclei or spindles analyzed per embryo. In B and E, statistical significance was evaluated with an unpaired t-test (in E, comparisons were between mean values per embryo). *, P<0.05; ns, not significant. Scale bars: A and C, 10 µm; D distance, 2 µm; D time, 30 s.

Dynein has also been implicated in creating tension between sister kinetochores in both mammals and flies (Siller *et al*, 2005; Varma *et al*, 2008; Yang *et al*, 2007). This process contributes to SAC inactivation in several systems, potentially by stabilizing kinetochore-microtubule attachments (Maresca & Salmon, 2010). To assess interkinetochore tension, we measured the distance between sister kinetochores using the centroids of Spc25-RFP signals.

This analysis revealed that, whilst there was no difference in the mean interkinetochore distance between wild-type and *S3372C* metaphase spindles (Figure 7B), there was substantially more variability in this metric in the mutant condition (Levene test, P = 0.001). These data raise the possibility that the S3372C mutation results in variable levels of tension between kinetochore pairs.

A particularly well-characterized mitotic role of dynein in silencing the SAC is transporting checkpoint proteins away from kinetochores along the attached microtubules (so-called ‘streaming’; Basto *et al*, 2004; Gassmann *et al*, 2010; Griffis *et al*, 2007; Hinchcliffe & Vaughan, 2017; Howell *et al*, 2001; Wojcik *et al*, 2001). Dynein co-operates in this process with Spindly, which interacts with the Rod-Zwilch-Zeste White 10 (RZZ complex) to link SAC components to the motor (Gama *et al*, 2017; Griffis *et al*, 2007; Mosalaganti *et al*, 2017). To assess the transport of SAC components in *S3372C* embryos, we monitored the behavior of GFP-tagged Rod using time-lapse imaging. GFP-Rod showed an abnormal build-up at the kinetochore in *S3372C* mutants (Figure 7C and D), consistent with the previously observed strong accumulation of dynein at this site (Figure 6A). However, streaming of GFP-Rod could be observed in the mutant embryos, albeit at a significantly lower velocity than in the wild-type condition (Figure 7C – E and Movie S4).

It is conceivable that alterations in any one of kinetochore-microtubule attachments, tension between sister kinetochores, and dynamics of Rod streaming (or a combination of these scenarios) are sufficient to prevent SAC inactivation in *S3372C* embryos and thereby cause metaphase arrest. To determine if the SAC might remain active in the mutant embryos, we examined the distribution of Cyclin B, which is degraded by APC/C once the checkpoint is satisfied (reviewed by Huang & Raff, 1999; Murray, 1995; Musacchio, 2015). Whereas Cyclin B was barely detectable in the vicinity of the spindle apparatus in late metaphase and anaphase in wild-type embryos, it was associated with metaphase-arrested *S3372C* spindles (Figure S6A). Persistence of Cyclin B on the mutant spindles was not due to a failure to recruit Cdc20/fzy – the co-activator of APC in early embryos – to its sites of enrichment at the centrosome and kinetochore (Raff *et al*, 2002) (Figure S6B). These data are consistent with the S3372C mutation blocking the metaphase-anaphase transition by preventing SAC silencing.

To test if this is actually the case, we introduced a null mutation in the gene encoding the checkpoint protein Mad2 into the *S3372C* background. Because the SAC is inactive in *Mad2* mutants (Buffin *et al*, 2007; Defachelles *et al*, 2015), the *S3372C* mitotic arrest should be rescued in the double mutants if it is primarily caused by a failure to alleviate SAC inhibition. However, this was not the case – whilst there was an increase in the proportion of spindles in telophase in *Mad2 S3372C* double mutant embryos (14.3% vs 0.8% in *S3372C* single mutant embryos; Fisher’s exact test, P <0.001), the vast majority of spindles still arrested at metaphase (Figure 7F and G). Although this result does not rule out the MTBD mutant impairing SAC silencing through the mechanisms described above, it suggests that this can make, at best, a minor contribution to the mitotic arrest phenotype. Thus, our data point to a novel, SAC-independent function of dynein that is required for the metaphase to anaphase transition in embryos.

### The S3372C mutation specifically affects dynein behavior under load

Next, we attempted to elucidate how the S3372C mutation selectively affects anaphase progression. We first considered the possibility that the mutation impairs the interaction of dynein with a specific tubulin isotype that is important for this process. *Drosophila* has four α-tubulin isotypes and four β-tubulin isotypes (Nielsen *et al*, 2010). One of these proteins, α4-tubulin (also known as α-tubulin67C or maternal α-tubulin), is only expressed during the first two hours of embryogenesis (Matthews *et al*, 1989) and is known to co-operate with the ubiquitously expressed α1-tubulin (also known as α-tubulin84B) to facilitate early nuclear divisions (Mathe *et al*, 1998; Matthews *et al*, 1993; Venkei *et al*, 2006). These observations make α4-tubulin the most likely tubulin isotype to mediate the specific effects of the S3372C mutation. Immunostaining revealed that whereas α1-tubulin was present throughout the spindle apparatus, α4-tubulin was enriched at the spindle poles (Figure S7A). Thus, α4-tubulin does not seem well placed to mediate the effects of S3372C dynein at the kinetochore. More significantly, although α4-tubulin and α1-tubulin have only 67% amino acid identity, the dynein-binding region of the two proteins is identical (Figure S7B). Thus, it seems unlikely that the S3372C mutation specifically affects the interaction of dynein with α4-tubulin.

These observations led us to explore the alternative idea that S3372C alters a general aspect of dynein behavior that is particularly important for anaphase progression in embryos. We therefore set out to determine the effects of the S-to-C mutation on the activity of purified dynein complexes *in vitro*. To this end, the equivalent mutation (S3386C) was introduced into the full human dynein complex, for which a recombinant expression system is available (Schlager *et al*, 2014; Figure S8A). As expected from its position in the MTBD, this mutation did not prevent the association of the heavy chain with the other dynein subunits (Figure S8B).

We first monitored the movement of tetramethylrhodamine (TMR)-labeled wild-type and S3386C mutant dynein complexes along immobilized pig brain-derived microtubules in the presence of dynactin and BICD2N using total internal reflection fluorescence (TIRF) microscopy (Figure 8A). It is in this assay that the human disease-associated mutations were found to significantly impair dynein activity (Hoang *et al*, 2017). In contrast, the S-to-C mutation had no effect on the frequency of processive movements of dynein, or the distance or velocity of these events (Figure 8B – D and Figure S8C and D). Thus, the novel MTBD mutation does not perturb the translocation of isolated dynein-dynactin-activating adaptor complexes along microtubules.

**Figure 8.**
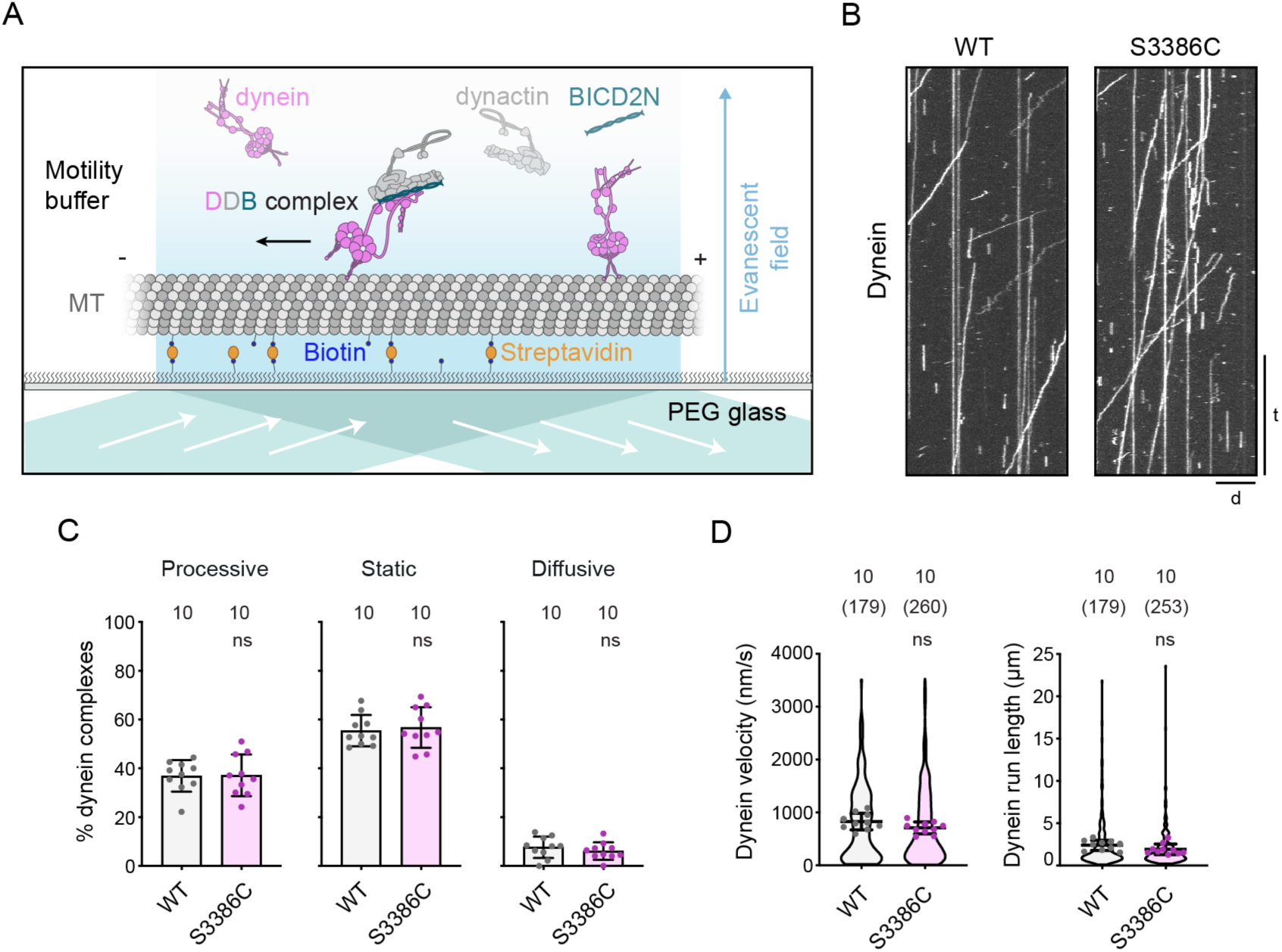
The S-to-C mutation does not alter motility of an isolated dynein-dynactin-activating adaptor complex. (A) Diagram of TIRF microscopy-based *in vitro* motility assay using an assembly of TMR-labeled dynein, dynactin and BICD2N. In the absence of dynactin and BICD2N, dynein is autoinhibited and not capable of long-range transport. MT, microtubule; PEG, polyethylene glycol (used for passivation). (B) Example kymographs of wild-type (WT) or S3886C TMR-ådynein motility in the presence of dynactin and BICD2N in the assembly mix. There were no consistent differences in the intensity of dynein signals between the wild type and mutant across the experiments. Microtubule minus end is to the left. d, distance; t, time. Scale bar: distance, 5 µm; time, 20 s. (C, D) Quantification of (C) percentage of microtubule-associated dynein complexes that exhibit processive, static or diffusive behavior and (D) velocity and run length of the processive dynein fraction. In C, columns display mean values for individual microtubules; error bars are S.D.; circles are values for individual microtubules. Numbers above each column indicate the total number of microtubules in each case. In D, violin plots show values for individual motile DDB complexes and circles show mean values per microtubule. Horizontal lines shown mean ± S.D. of values for individual DDB complexes. Numbers without parentheses above bars are number of microtubules, with total numbers of motile DDB complexes given in parentheses. In C and D, statistical significance was evaluated with an unpaired t-test (in D, comparisons were between mean values per microtubule). ns, not significant. Data were summed from 2 experiments per condition. See Figure S8C and D for histograms of velocity and run length.

The absence of cargo in the above assays meant that dynein was operating under minimal load. To determine if the S-to-C mutation influences dynein performance under resistive loads, we quantified force production by the motor using high-resolution optical trapping (Figure 9A). Polystyrene beads were sparsely decorated with BICD2N-GFP via a GFP antibody, incubated with dynactin and either wild-type or S3386C dynein, and brought near a surface-immobilized microtubule using an optical trap (Belyy *et al*, 2016). Fixing the position of the trap while the motor carries the bead along the microtubule allows quantification of the force at which the bead detaches from the microtubule after a brief stall (referred to as the stall force). Whilst performing these experiments we noticed that, whereas the wild-type DDB complex moved rapidly along the microtubule at low resistive forces (∼1 pN), the S3386C version had a tendency to pause in these conditions (Figure 9A; e.g. green arrowheads). Following these pauses, the mutant DDB either detached from the microtubule or continued to move forward. These data indicated that S3386C dynein is sensitive to external load. Confirming this notion, whereas the stall force of beads driven by wild-type DDB was 4.3 ± 0.1 pN (mean ± S.E.M.), consistent with previous observations (Belyy *et al*, 2016), the value was reduced to 2.9 ± 0.1 pN for the mutant (Figure 9B). Thus, the S-to-C mutation reduces dynein’s peak force production by one-third. The mutant complexes also had longer stall times than the wild type (Figure 9C), which may be related to the stalling of the motor at lower resistive forces. These results show that mutant dynein exhibits defective motility in the presence of external load.

**Figure 9.**
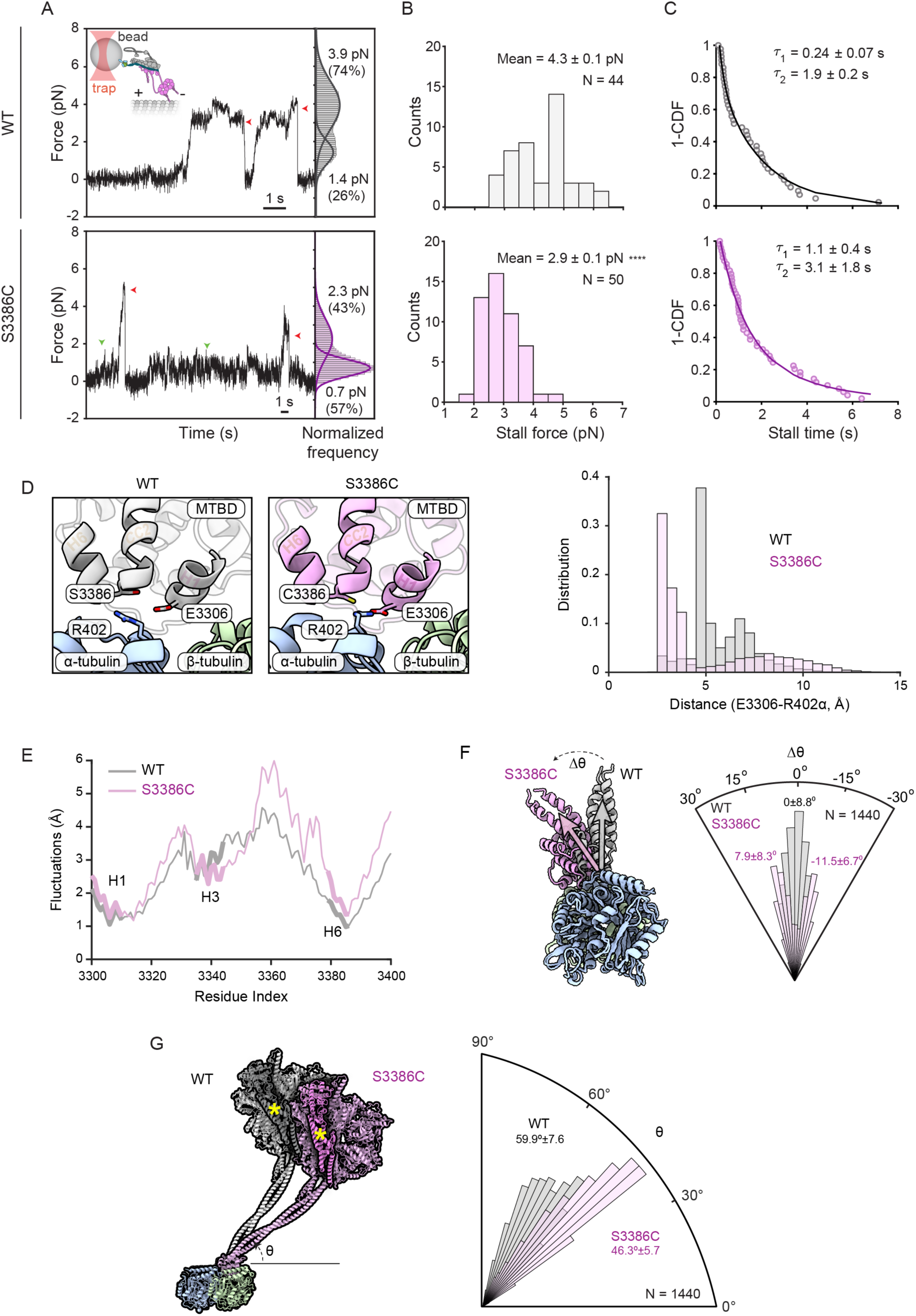
The S-to-C mutation alters dynein-dynactin-activating adaptor behavior under load and the position of the dynein MTBD and stalk relative to the microtubule. (A) Left, representative traces of beads driven by a single wild-type (WT) or S3386 DDB complex in a fixed optical trapping assay. Traces were downsampled to 250 Hz. Red arrowheads show detachment of the motor from a microtubule after the stall; green arrowheads show pausing of the mutant dynein at low forces. Inset cartoon shows how DDB is attached to beads from its BICD2N subunit via GFP-antibody linkage. Right, normalized histograms showing the probability of a dynein-driven bead sampling different forces under the trap. Dwelling of the beads at 0 pN force when they are not tethered to the microtubule was subtracted from the histogram. Solid curves represent a multiple Gaussian fit to calculate the mean and percent population of the 2 peaks. (B) Stall forces of DDB complexes with WT and mutant dynein (errors are S.E.M.; N is number of stalls). Statistical significance was evaluated with an unpaired t-test. ****, P<0.0001. (C) Inverse cumulative distribution (1-CDF) of motor stall times. Time constants (*τ* ± S.E.M.) were calculated from a fit to a double exponential decay (solid curves). In A – C, data are from 11 beads in 5 independent experiments for WT and from 14 beads in 7 independent experiments for mutant. (D) Left, example snapshots of E3306(dynein) and R402(α-tubulin) positioning in MD simulations of the WT and S3386C human dynein MTBD plus a portion of the stalk in the presence of the α/β-tubulin dimer. Right, distance distributions of E3306(dynein)-R402(α-tubulin) in WT and S3386C dynein simulations. (E) Fluctuations of WT and S3386C MTBD and portion of the stalk with respect to the microtubule in MD simulations. H1, H3, and H6 helices are highlighted with thicker lines. (F) Left, superimposition of examples from simulations with the WT and S3386C MTBD plus a portion of stalk when viewed down the longitudinal axis of the microtubule. Gray and magenta vectors show most frequent MTBD orientation for WT and S3386C, respectively. Right, angular orientation histogram (normalized frequency) for the MTBD around the longitudinal axis of the microtubule in simulations. The WT distribution exhibited a single peak (assigned a value of 0), whereas the S3386C mutant exhibited a double-peaked distribution (peaks of fitted curves ± S.D. are shown). (G) Left, superimposition of examples from simulations with WT and S3386C MTBD plus a portion of stalk when viewed facing the longitudinal axis of the microtubule. The remainder of the dynein motor domain structure (PDB, 7Z8F; Chaaban & Carter, 2022) was superimposed on the simulations to show predicted position of the linker (darker shading and yellow asterisk) and AAA+ ring. Right, angular orientation histogram (normalized frequency) for the stalk relative to the longitudinal axis of the microtubule in simulations (peaks of distributions ± S.D. are shown). In F and G, N is number of trajectories for each type of complex.

### S3372C modulates dynein’s microtubule interaction and stalk positioning

Remarkably, the effects of the S-to-C change on dynein’s function in embryonic mitosis and its behavior under load stem from the substitution of a single oxygen atom for a sulfur atom within the ∼90,000-atom Dhc polypeptide. To evaluate the effects of the S-to-C change on the structure and dynamics of the MTBD, as well as its interaction with the microtubule, we performed all-atom MD simulations of the human wild-type and S3386C MTBD together with a portion of the stalk bound to the α/β-tubulin dimer (modeled on the cryo-EM structure of Lacey *et al*, 2019). Four sets of 900-ns-long simulations were performed for each variant in the presence of explicit solvent.

Analysis of the ensemble of conformations in the simulation trajectories of the wild type revealed that S3386 did not make contact with the α/β-tubulin dimer. The residue did, however, frequently form hydrogen bonds with the main chain of residues in H6 of the MTBD (V3382) and the base of CC2 of the stalk (C3389) (Figure S8E and Table S1). Thus, the serine appears to stabilise the interaction of H6 with the stalk. In the simulations with the mutant dynein, the overall fold of the MTBD was not overtly different from the wild type. C3386 also did not make contact with tubulin and had a very similar frequency of interactions with V3382 and C3389 to that observed with S3386 (Figure S8E and Table S2). However, the mutation led to hydrophobic interactions with several neighboring residues in the MTBD that were not evident in the wild type (Figure S8E and Table S3). This was accompanied by frequent formation of a salt bridge between E3306 within H1 of the MTBD and R402 of α-tubulin, which was seldom seen in the wild-type simulations (Figure 9D; frequency of 62.4% and 8.4% in mutant and wild-type simulations, respectively). The mutation also increased the magnitude of fluctuations of several elements of the MTBD relative to the microtubule (mean of 16.9% compared to the wild type), with those located between the end of H3 and the start of CC2 of the stalk showing the greatest effect (mean of 25.7% compared to the wild type) (Figure 9E). Thus, the S-to-C mutation alters the MTBD’s interaction with the microtubule, including by increasing the relative flexibility of the MTBD.

Further analysis demonstrated that, whilst the registry of the stalk was not sensitive to the S-to-C change, the mutation induced either clockwise or anticlockwise tilting of the MTBD and base of the stalk around the longitudinal axis of the α/β-tubulin dimer (Figure 9F; peak of angle distribution for the mutant 11.5° clockwise and 7.9° anticlockwise compared to the wild type). In addition, the angle of the stalk relative to the longitudinal axis of the tubulin dimer was decreased by the mutation (Figure 9G; peak of angle distribution 59.9° and 46.3° for wild-type and S3386C dynein, respectively). As discussed below, these changes in stalk positioning relative to the microtubule could account for the force-sensitive nature of the mutant motor complex.

## DISCUSSION

### Mutational analysis of Dhc *in vivo*

The diverse cellular roles of dynein can complicate functional analysis of the motor significantly. We set out to address this problem by combining mutagenesis in *Drosophila* with *in vitro* analysis of motor activity. We first assessed the phenotypes of disease-associated missense mutations in Dhc for which *in vitro* effects have already been defined. Abnormalities were only manifest in homozygous mutant *Drosophila*, whereas the equivalent mutations in human or mouse cause neurological disease when heterozygous (Poirier *et al*, 2013; Schiavo *et al*, 2013; Vissers *et al*, 2010; Willemsen *et al*, 2012). This may reflect the relatively short length of *Drosophila* neurons making them less sensitive to partially impaired cargo transport. Nonetheless, the homozygous *Drosophila* phenotypes provide valuable information about the consequences of *Dhc* disease-associated mutations in an animal model, and thereby extend previous work on these lesions *in vitro* (Hoang *et al*, 2017) and in budding yeast (Marzo *et al*, 2019). For example, our observation that disease-linked mutations can cause abnormal accumulation of synaptic vesicles and impaired retrograde transport of mitochondria in axons supports our hypothesis that they interfere with the motility of cargo-motor complexes in neurons (Hoang *et al*, 2017).

The scope of our study was increased by the recovery of novel *Dhc* alleles whilst generating the disease-associated mutations. Collectively, the phenotypic characterization of the disease-associated and novel mutations establishes an allelic series that will facilitate studies of dynein’s involvement in multiple processes during development and in adulthood. Some of the mutations we investigated cause phenotypes – synaptic vesicle accumulations, impaired mitochondrial transport, and short bristles – that were previously seen with classical hypomorphic mutations in *Dhc* or when the function of the gene is knocked down with RNAi (Gepner *et al*, 1996; Martin *et al*, 1999; Melkov *et al*, 2016; Vagnoni *et al*, 2016). Thus, these mutations may impair core dynein functions to varying degrees, rather than affect discrete processes. Our earlier finding that the disease-associated mutations reduce the motility of purified dynein-dynactin-activating adaptor complexes to different extents (Hoang *et al*, 2017) is consistent with this scenario.

In sharp contrast, the novel MTBD mutation, S3372C, only affects a subset of dynein functions. This allele does not cause visible abnormalities in adults, yet homozygous mothers are infertile because mitotic spindles in their embryos arrest at metaphase. Whilst some classical hypomorphic dynein alleles, as well as the F597Y mutation, also have maternal effects on embryogenesis, these only become evident at later embryonic stages and are accompanied by morphological phenotypes in the parents (Gepner *et al*, 1996; Melkov *et al*, 2016; Robinson *et al*, 1999; Wilkie & Davis, 2001; this study). Consistent with the normal appearance of *S3372C* adults, we found that the mutation does not impair dynein-dependent functions during mitosis in larval neuroblasts, or mitochondrial transport in axons of adult neurons. Collectively, these analyses demonstrate that S3372C disrupts a specific, maternal activity of dynein that is critical for early embryogenesis. The effect of S3372C contrasts sharply with that of the neighbouring R3370Q mutation, which causes zygotic mutants to arrest during larval stages. These findings validate the use of mutagenesis in flies to dissect the *in vivo* functions of the motor complex.

### The mechanistic basis of S3372C’s selective phenotype and force sensitivity

The observations described above raised the question of how the S3372C mutation has such a selective effect on Dhc activity. The finding that embryos of heterozygous *S3372C* mothers have no developmental defects is consistent with the allele causing a loss, rather than a gain, of dynein function. We have shown that neither stage-specific instability of Dhc protein nor ectopic disulfide bonding within it is responsible for the embryo-specific phenotype. We have also provided evidence that an altered interaction of dynein with the maternal α4-tubulin isotype is not a contributing factor. Whilst we cannot rule out the S-to-C change blocking a phosphorylation event on S3372 that is important for anaphase onset, this seems unlikely because the frequent hydrogen bonding of this sidechain with neighboring residues in H6 and CC2 means it should not be readily accessible to a kinase. Indeed, there is no evidence that S3372 is post-translationally modified in previous proteome-wide studies of embryos from multiple *Drosophila* species (Hu *et al*, 2019) or our ongoing analysis of dynein complexes immunoprecipitated from *D. melanogaster* embryos.

However, using high-resolution optical trapping, we observed a striking effect of the S-to-C mutation on the behavior of dynein-dynactin-activating adaptor complexes operating with a resistive load. Whereas motility was normal in the absence of load, mutant complexes exhibited excessive pausing when exposed to a resistive force, as well as a substantially reduced peak stall force. Therefore, the most parsimonious explanation for the restricted *in vivo* phenotype is that the metaphase to anaphase transition in the embryo needs dynein to work in a specific load regime that is problematic for the mutant motor complex. Consistent with this notion, we observed a reduced velocity of dynein-driven motion of GFP-Rod away from the metaphase plate in *S3372C* mutant embryos.

Our MD simulations of the human Dynein heavy chain’s interaction with the microtubule allow us to speculate how the mutation causes force sensitivity. Our data indicate that replacement of serine’s hydroxyl group with the more hydrophobic sulfhydryl group of cysteine (Catalano *et al*, 2021) causes several new hydrophobic interactions with neighbouring residues, which are accompanied by increased flexibility of the MTBD, as well as repositioning of the stalk, relative to the microtubule. Our analyses suggest that the MTBD and stalk in the mutant dynein tilt abnormally in both the anticlockwise and clockwise direction in the plane perpendicular to the longitudinal axis of the microtubule. Thus, the stalk appears to be more mobile in this plane in the mutant, which may reduce dynein’s ability to overcome an opposing force. The mutation also reduced the angle between the stalk and the longitudinal axis of the microtubule, a change that is expected to increase the distance the linker moves with respect to this axis during its powerstroke. This would mean that more work needs to be done against the opposing horizontal force during the linker swing, which could contribute to the reduced force output. An interesting implication of these ideas for wild-type dynein mechanism is that the MTBD plays an important role in maximising motor performance under load by controlling stalk orientation.

### Insights into dynein function in mitosis

As described in the Introduction, a prime example of the difficulties of disentangling dynein’s *in vivo* functions is in mitosis, where the motor has been implicated in a wide range of processes. Partial knockdowns with RNAi, function-blocking antibodies and small molecule antagonists have each been used to circumvent the cell lethality that results from complete loss of dynein activity. Unfortunately, each of these approaches simultaneously targets many aspects of motor function, which can indirectly affect the specific mitotic process being studied. In the *Drosophila* embryo, dynein’s roles in mitosis have also been investigated with hypomorphic mutations (Robinson *et al*, 1999). However, in addition to being likely to affect multiple dynein-dependent events, these mutations have weak mitotic phenotypes since sufficient protein function must remain to produce viable mothers. Our discovery of a missense mutation that strongly affects nuclear divisions in the embryo without disrupting other dynein functions offers a unique tool to study the mitotic roles of the motor.

Although we cannot exclude an indirect effect of the S3372C mutation on dynein’s functions in other parts of the spindle apparatus, such as the poles and centrosomes, the build-up of dynein at the kinetochore, as well as slower transport of Rod away from this site, suggest that impaired kinetochore functions of dynein make an important contribution to the metaphase arrest. In light of our discovery of force sensitivity imparted by the S-to-C mutation, we can speculate that tight bundling of microtubules at kinetochores in embryonic spindles, or other physical constraints of this environment, provide a strong opposition to motility that cannot be overcome by the mutant motor. In such a scenario, the ectopic build-up of S3372C dynein in the vicinity of the kinetochore and associated microtubules may be the manifestation of the increased pausing of the motor complex observed *in vitro* under resistive loads. According to this view, the failure of S3372C to block mitosis in other cell types, include L3 neuroblasts, could reflect differences in the forces encountered by dynein near the kinetochore. Alternatively, there may be redundant mechanisms for initiating anaphase in these systems.

Remarkably, the MTBD mutation does not appear to block anaphase progression in embryos by preventing the well-characterized role of kinetochore-associated dynein in silencing the SAC, as the defect persists when the checkpoint is inactivated by mutation of Mad2. Collectively, these observations indicate that kinetochore dynein has a novel role in licensing the transition from metaphase to anaphase. We found that this function is not associated with a failure to localize the APC/C co-activator Cdc20/fzy to the spindle apparatus. Dynein may therefore directly promote coupling of Cdc20/fzy to APC/C. Alternatively, APC/C activation may be triggered indirectly by another kinetochore-associated process that depends on the motor. For example, the apparent variability in tension between sister kinetochores in *S3372C* embryos, which could reflect abnormal force generation by the mutant motor complex, might prevent APC/C activation through the complex series of signalling events that respond to chromosome biorientation (Fujimitsu & Yamano, 2021; Krenn & Musacchio, 2015; Liu *et al*, 2012; McVey *et al*, 2021). It is also possible that force production by dynein plays a physical role in separating sister chromosomes downstream of APC/C activation, or that the ectopic accumulation of dynein and associated proteins at the kinetochore in the mutant embryos impairs engagement of other important factors with this structure. Investigating these, as well as other, potential explanations for the metaphase block in the *Drosophila* embryo will be the goal of future studies.

### Outlook

As well as pointing to novel mechanisms controlling anaphase progression, the S-to-C mutation in the MTBD may be valuable in other contexts. As the mutated serine is widely conserved, including in dynein-2 and axonemal dyneins, its substitution with cysteine may allow load-dependent functions of dynein family members to be dissected in other contexts. Moreover, if we are correct and the mutation causes force-responsive dwelling of the motor on microtubules *in vivo*, the location of the mutant dynein complexes may act as a reporter of subcellular regions and events where the motors are experiencing high load. This information could be useful for producing quantitative models of motor behavior *in vivo*. We also anticipate that our results will stimulate further efforts to dissect the function of cytoskeletal motors by genetic manipulation of specific mechanical properties. Whilst *Drosophila* is an attractive organism in which to pursue this work because of the ease of gene editing and ability to study motor function in cells within tissues, the same approach can of course be taken in other systems.

## AUTHOR CONTRIBUTIONS

D.S.G. and S.L.B. conceived the project. D.S.G., L.J., A.H., M. Golcuk, E.G., S.C., F.P., V.J.P.H and S.L.B. performed experiments and analyses. A.V., M.A.M., E.D., R.G., A.P.C, M. Gur, A.Y and S.L.B. provided training and supervision. S.L.B. co-ordinated the project. D.S.G. and S.L.B. drafted the manuscript, which was edited and approved by the other authors.

## ACKNOWLEDGMENTS

We thank members of the Bullock group and MRC-LMB fly community for support and input, as well as many members of the broader *Drosophila* community for reagents and discussions. We also thank MRC-LMB’s VisLab for help with illustrations and movies. The work in S.L.B.’s, E.D.’s, and A.P.C.’s groups was supported by the Medical Research Council, as part of United Kingdom Research and Innovation (file reference numbers MC_U105178790 (S.L.B.), MC_UP_1201/13 (E.D.) and MC_UP_A025_1011 (A.P.C)). The work was also supported by a NC3Rs David Sainsbury Fellowship (to A.V), a Marie-Curie IntraEuropean Fellowship (to F.P.), and awards from NIH (GM136414) and NSF (MCB-1055017 and MCB-1617028) (to A.Y.), the Fondation pour la Recherche Médicale Equipe Labellisée (DEQ20170336742) (to R.G), an EMBO Postdoctoral Fellowship (ALTF 334-2020) (to S.C.), the PRACE-Partnership for Advanced Computing in Europe (PRA205144 and PRA2021250119) and the Scientific and Technological Research Council of Türkiye (TUBITAK, 121C283) (to M. Gur). M.A.M. is supported by a BBSRC project grant (BB/T00696X/1) (to S.L.B.). For the purpose of open access, the MRC Laboratory of Molecular Biology has applied a CC BY public copyright license to any Author Accepted Manuscript version arising.

## MATERIALS AND METHODS

### *Drosophila* culture and existing strains

*Drosophila* strains were cultured on Iberian fly food (5.5% [w/v] glucose, 5% [w/v] baker’s yeast, 3.5% [w/v] organic wheat flour, 0.75% [w/v] agar, 0.004% [v/v] propionic acid, 16.4 mM methyl-4-hydroxybenzoate [Nipagin]) at 25°C and 50% ± 5% relative humidity with a 12h-light/12h-dark cycle. The following previously established strains and alleles were used in this study: *w^1118^* (Bullock lab stocks); *yw* (Bullock lab stocks); *Dhc^null^* (Fumagalli *et al*, 2021); *appl-* Gal4 (Torroja *et al*, 1999); *UAS-mito::GFP* (Pilling *et al*, 2006); *insc-Gal4>UAS-ChRFP::α-Tubulin* (Gallaud *et al*, 2022; Hartenstein *et al*, 2015); *His2Av::mRFP* (Heeger *et al*, 2005); *Jupiter::GFP* (Morin *et al*, 2001); *Asl::mCherry* (Conduit *et al*, 2015); *Spc25::mRFP* (Schittenhelm *et al*, 2007); *GFP::Dlic* (Pandey *et al*, 2007); *Dhc::3HA* (*Dhc* genomic rescue construct; Iyadurai *et al*, 1999); *GFP::Rod* (Basto *et al*, 2004); *GFP::fzy* (Raff *et al*, 2002); and *mad2^P^* (Buffin *et al*, 2007). The *mad2^P^ S3372C* strain was generated by recombination. The presence of the *mad2* null allele in the *mad2^P^* and *mad2^P^ S3372C* strains was confirmed by PCR and Sanger sequencing.

### CRISPR/Cas9-mediated knock-in of *Dhc* mutations

Mutations in the *Drosophila Dhc64C* gene were generated using previously established procedures (Port & Bullock, 2016; Port *et al*, 2014). Briefly, pCFD3 plasmids were generated that express, under the control of the U6:3 promoter, single gRNAs that target *Dhc* close to the codon to be mutated (see Table S4 for sequences of oligonucleotides used for gRNA cloning). For the experiments designed to produce F579Y, R1951C, R3370Q, S3372C + C3375S, C3375S and H3808P, transgenic strains expressing the gRNAs were established, with males of these strains crossed to *nos-cas9* females (CFD2 strain; Port *et al*, 2014) to produce *nos-cas9/+; gRNA/+* embryos. These embryos were injected with a 500 ng μl^-1^ solution of a donor oligo (Ultramer, IDT) that codes for the desired missense mutation (see Table S4 for donor sequences). In cases where the mutation would not disrupt targeting by the gRNA, synonymous changes that prevent recutting of the modified allele by the Cas9/gRNA complex were also introduced (Table S4). For the generation of the other mutations, the above procedures were replicated, except the gRNAs were introduced by co-injection of the pCFD3-gRNA plasmid (100 ng μl^-1^ solution) with the donor oligo into *nos-cas9* embryos.

Flies containing the desired mutation were identified by sequencing of PCR products containing the target region, as described (Port & Bullock, 2016), with stocks established using balancer chromosomes. Other in-frame mutations, which resulted from imprecise repair of Cas9-mediated DNA cleavage, were also retained for phenotypic analysis. With the exception of *S3372C* and *R3370Q*, all mutations were isogenized by backcrossing to the *w^1118^* strain for 6 – 10 generations. In addition, we balanced a chromosome in which *Dhc* had not been mutated during the CRISPR process. This ‘CRISPR WT’ strain was used as the control genotype for a subset of experiments.

### Assaying lethality and fertility

To assess lethality, flies heterozygous for the *Dhc* missense mutations (or the CRISPR WT chromosome) and balanced with TM6B were crossed together or with *Dhc^null^/TM6B* flies. Absence of non-TM6B adult offspring indicated developmental arrest of homozygotes or *trans*-heterozygotes. To assess the stage of development arrest, crosses were performed with stocks balanced with the fluorescent balancer *TM3 [actin5C>GFP]*. In these experiments, cohorts of homozygous (GFP-negative) embryos were transferred to plates containing apple-juice agar (1.66% [w/v] sucrose, 3.33% [w/v] agar, 33.33% [v/v] apple juice and 10.8 mM methyl-4-hydroxybenzoate), and the number of animals that survived until early L2, late L2, L3 and pupal stages scored through regular inspections. Genotypes that did not arrest before pupal stages but did not reach adulthood were classed as pupal lethal and this was confirmed in independent crosses with the TM6B balancer, which has the *Tb* marker that is visible at pupal stages.

Fertility of *Dhc* mutant and control females was assessed in crosses to wild-type (*w^1118^*) males 5 days after eclosion of the females. After 24 h, crosses were transferred to egg-laying cages capped with apple-juice agar plates and the proportion of total embryos that had hatched 28 – 48 h after egg-laying was recorded in multiple technical replicates over the next 3 days.

### Immunostaining

L3 wandering larvae were dissected and fixed in 4% formaldehyde as described (Hurd & Saxton, 1996). Embryos were collected, dechorionated, fixed at the interface of 4% formaldehyde and n-heptane, and devitellinized using standard procedures (Port *et al*, 2014). Larval preparations were washed in PBS/0.1% Tween (PBT) and blocked in 20% Western Blocking Buffer (Sigma Aldrich) in PBT, whereas embryos were washed in PBS/0.1% Triton X-100 (PBST) and blocked in 20% Western Blocking Buffer in PBST. Details of primary and secondary antibodies, including working dilutions, are provided in Tables S5 and S6. Samples were mounted in Vectashield containing DAPI (Vector Laboratories). Segmental nerves were imaged with a Zeiss 780 laser-scanning confocal microscope using a 40x/1.3 NA oil-immersion objective. Embryos were typically imaged with a Zeiss 710 or 780 laser-scanning confocal microscope using a 63x/1.4 NA oil-immersion objective, the exception being the use of a Nikon T2 wide-field microscope equipped with a 20x/0.75 NA air objective for the initial documentation of the stage of embryonic arrest of *S3372C* mutants.

### Assessing mitochondrial transport in the wing nerve

Fly wing mounting, imaging and analysis were performed as described previously (Vagnoni & Bullock, 2016; Vagnoni *et al*, 2016). Briefly, CO_2_-anethetized male or female flies that had eclosed 2 days earlier were mounted on double-sided sticky tape and wings coated in Voltalef 10S halocarbon oil (VWR). Movements of mitochondria (labeled with mito-GFP under the control of the neuronal driver *appl-GAL4*) in the arch region of the wing nerve were visualized with an UltraVIEW ERS spinning disk system (PerkinElmer) equipped with an Orca ER Charge-coupled device (CCD) camera (Hamamatsu) using a 60x/1.2 NA oil-immersion objective on an IX71 microscope (Olympus). A single focal plane was imaged with an acquisition rate of 0.5 frames s^-1^ for 3 min. Images were processed and analyzed with ImageJ (Schneider *et al*, 2012). The genotypes of the image series were hidden from the experimenter using the BlindAnalysis macro (Steve Royle, University of Warwick), followed by image straightening, stabilization and manual tracking of motile mitochondria in MTrackJ, as described previously (Vagnoni & Bullock, 2016; Vagnoni *et al*, 2016). We additionally quantified the total number of mitochondria per 50 μm of the wing nerve and the proportion of these that underwent transport (i.e. had at least one continuous bout of net motion of ≥ 2 μm [a ‘run’]).

### Analysis of *Drosophila* neuroblast divisions

Brains of L3 larvae (∼120 h after egg laying) were dissected in Schneider’s media (Sigma Aldrich) containing 10% foetal calf serum (ThermoFisher Scientific) and transferred to 50 µl wells of Ibidi Angiogenesis µ-Slides for live imaging. Mutant and control brains were imaged in parallel at 25°C. Z-series with a height of 20 µm and 1-µm spacing were acquired every 30 s using a spinning disk system consisting of a Leica DMi8 microscope equipped with a 63x/1.4 NA oil-immersion objective, a CSU-X1 spinning disk unit (Yokogawa) and an Evolve Electron-Multiplying CCD (EMCCD) camera (Photometrics). The microscope was controlled by Inscoper Imaging Suite software (Inscoper). Images were processed with Fiji (Schindelin *et al*, 2012).

### SDS-PAGE and immunoblotting

*Drosophila* embryo extracts were generated for immunoblotting as described (McClintock *et al*, 2018). SDS-PAGE and protein transfer to polyvinylidene difluoride (PVDF) membrane (Immobilon-P, Merck Millipore) was performed with the NuPAGE Novex and XCell II Blot Module systems (ThermoFisher Scientific) according to the manufacturer’s instructions. Membranes were blocked with 5% dried skimmed milk powder (Marvel) and washed in PBS/0.05% Tween-20. Details of primary and secondary antibodies, including working dilutions, are provided in Tables S5 and S6. Secondary antibodies were detected with ECL Prime reagents (Cytiva Amersham) as instructed by the manufacturer.

### Sequence and structure analysis

Alignments of Dynein heavy chain and α-tubulin sequences were produced with ESPript 3.0 (Robert & Gouet, 2014; https://espript.ibcp.fr/ESPript/ESPript/). Visualization and analysis of experimentally determined MTBD structures was performed with PyMOL (version 2.5.1; Schrödinger) and ChimeraX 1.2.5 (Goddard *et* al, 2018; https://www.cgl.ucsf.edu/chimerax/). The cryo-EM structure of the mouse dynein-1 MTBD and portion of the stalk bound to the microtubule (PDB 6RZB; Lacey *et al*, 2019) was generated from a ‘cysteine-light’ dynein in which C3389 was mutated to alanine. The alanine residue was therefore substituted for the native cysteine in Figure 3B.

Structure predictions were performed using a local installation of ColabFold 1.2.0 (Mirdita *et al*, 2022), running MMseqs2 (Mirdita *et al*, 2019) for homology searches and Alphafold2 (Jumper *et al*, 2021) for predictions with 3 recycles. For the MTBD and stalk regions of different dyneins, predictions were performed with the following amino acid sequences: *D. melanogaster* Dhc (Uniprot ID P37276) 3212-3451, *H. sapiens* DYNC1H1 (Uniprot ID Q14204) 3227-3465, *H. sapiens* DYNC1H2 (Uniprot ID Q8NCM8) 2922-3162, and *H. sapiens* DYH7 (Uniprot ID Q8WXX0) 2614-2866. The top-ranking models were visualized in ChimeraX 1.2.5., with secondary structures colored based on their structural homology to mouse dynein (PDB 6RZB; Lacey *et al*, 2019).

### Live imaging of mitosis in *Drosophila* embryos

Dechorionated transgenic embryos with fluorescently-marked centrosomes, histones, microtubules, kinetochores, Dlic, Rod, or Fzy/Cdc20 were filmed under Voltalef 10S halocarbon oil (VWR) using the Ultraview ERS spinning disk system described above with a 60x/1.2 NA water-immersion objective or a Zeiss 710 laser scanning confocal with a 63x/1.4 NA oil-immersion objective. A single focal plane was imaged with an acquisition rate of 0.2 frames s^-1^ for up to 15 min to capture complete nuclear division cycles. To capture Rod streaming, an acquisition rate of 0.5 frames s^-1^ was used. Exposure times were typically maintained at 300 ms or 500 ms for microtubules, histones, Dlic and Rod, and at 1000 ms or 1500 ms for kinetochores and centrosomes. Images were processed with Fiji (Schindelin *et al*, 2012), including analysis of streaming of faint Rod signals with kymographs.

### Introduction of a C3386 codon into human DYNC1H1

Phusion High Fidelity Master Mix with GC buffer (New England Biolabs) was used for site-directed mutagenesis of pDyn1, which contains human DYNC1H1 sequences (accession number NM_001376.4) that are codon optimized for Sf9 insect cell expression. The presence of the mutation encoding the S3386C substitution, as well as the absence of other non-synonymous mutations, was confirmed using Sanger sequencing (Genewiz) and whole-plasmid next generation sequencing (MGH Center for Computational and Integrative Biology DNA Core).

### Production of dynein, dynactin and BICD2N

Wild-type and S3386C-containing human dynein complexes were produced as described (Hoang *et al*, 2017; Schlager *et al*, 2014). Briefly, they were expressed recombinantly in Sf9 cells by transposition into the baculovirus genome of sequences encoding wild-type or S3386 DYNC1H1 (tagged with SNAP for fluorescent labelling and ZZ (a synthetic Fc region-binding domain of protein A) for protein purification), as well as the other dynein subunits (DYNC1I2 [DIC2; AF134477], DYNC1LI2 [DLIC2; NM_006141.2], DYNLT1 [Tctex1; NM_006519.2], DYNLL1 [LC8; NM_003746.2] and DYNLRB1 [Robl1; NM_014183.3]). Dynein complexes were captured from Sf9 cell lysates using IgG Sepharose 6 FastFlow beads (GE Healthcare), labeled with SNAP-Cell-TMR-Star, and eluted with Tobacco Etch Virus protease by virtue of a cleavage site between the ZZ and SNAP tags. Dynein complexes were further purified by fast protein liquid chromatography (TSKgel G4000SWxl column [TOSOH Bioscience]) and concentrated by centrifugation through an Amicon Ultra-4 Centrifugal Filter Device (Merck Millipore). Native dynactin was purified from fresh pig brains as previously described (Schlager *et al*, 2014; Urnavicius *et al*, 2015) using a series of chromatography steps (XK 50/30 cationic exchange column [GE Healthcare] packed with SP-Sepharose Fast Flow [GE Healthcare], MonoQ HR 16/10 anionic exchange column [GE Healthcare] and TSKgel G4000SWxl column). SNAP-BICD2N and BICD2N-GFP (both containing residues 1-400 of mouse BICD2) were expressed and purified using the Sf9 baculovirus system, as previously described (Belyy *et al*, 2016; Schlager *et al*, 2014).

### Assessing motility of isolated dynein-dynactin-BICD2N complexes

Motility assays were performed using established protocols (Hoang *et al*, 2017; McClintock *et al*, 2018; Schlager *et al*, 2014). Briefly, biotin-and HiLyte-488-labeled pig brain microtubules were polymerized *in vitro* using commercial tubulin sources (Cytoskeleton Inc.) and stabilized with taxol (Sigma Aldrich) and GMPCPP (Jena Bioscience) before being adhered to a streptavidin-coated coverslip within the flow chamber. Approximately 30 min before imaging, a dynein-dynactin-BICD2N (DDB) assembly mix was prepared on ice by combining 100 nM TMR-labeled human dynein (wild type or S3386C), 200 nM pig dynactin and 1 µM SNAP-BICD2N in motility buffer (30 mM HEPES pH 7.3, 5 mM MgSO_4_, 1 mM EGTA pH 7.3, 1 mM DTT). A ‘motility mix’ comprising 2.5% (v/v) DDB assembly mix, 2.5 mM MgATP (Sigma Aldrich), 10% (v/v) oxygen scavenging system (1.25 µM glucose oxidase [Sigma Aldrich], 140 nM catalase [Sigma Aldrich], 71 mM β-mercaptoethanol and 25 mM D-glucose) in motility buffer supplemented with 50 mM KCl, 1 mg ml^-1^ α-casein and 20 µM taxol was introduced into the microtubule-containing flow chambers.

TIRF imaging was performed at room temperature (∼23 ± 1°C) using a Nikon TIRF system equipped with a back illuminated EMCCD camera iXon^EM+^ DU-897E (Andor) and a 100x/1.49 NA oil-immersion APO TIRF objective, and controlled with µManager software (Edelstein *et al*, 2010). Three movies were acquired per chamber, with acquisition for 500 frames at the maximum achievable frame rate (∼2 frames s^-1^) and 100-ms exposure per frame. Pixel size for each image series was 105 nm x 105 nm.

Analyses of dynein behavior was performed manually using kymographs in Fiji, as described (Hoang *et al*, 2017). Kymographs were generated from 5 microtubules per movie, followed by tracking of dynein movements with the Multipoint tool. Microtubule binding events were only counting if they lasted for ≥ 1.5 s (3 pixels in the y-axis) and processive events only counted when travel distance was ≥ 500 nm (5 pixels in the x-axis). Because DDB complexes could change speed during runs, mean velocity was calculated from individual velocity segments, as described previously (Schlager *et al*, 2014). The identities of kymographs were blinded before analysis using the BlindAnalysis macro in ImageJ.

### Optical trap-based force measurements

Optical trapping of DDB complexes was performed as described previously (Belyy *et al*, 2016). Briefly, DDB complexes were assembled with 1 μl of 0.38 mg ml^−1^ (mutant) or 1 μl 1.06 mg ml^-1^ (wild-type) dynein, 1 μl of 2.25 mg ml^−1^ dynactin, and 1 μl of 1.04 mg ml^−1^ BICD2N-GFP (Belyy *et al*, 2016) for 10 min at 4°C. The protein mixture was then added to 800-nm diameter carboxylated polystyrene beads that were coated with a rabbit polyclonal anti-GFP antibody (BioLegend MMS-118P), and incubated for 10 min. Flow chambers were decorated with sea urchin axonemes in MB buffer (30 mM HEPES pH 7.0, 5 mM MgSO_4_, 1 mM EGTA, 10% glycerol). The motor-bead mixture was then introduced to the chamber in imaging buffer (MB supplemented with oxygen scavenging system and 2 mM MgATP). The bead concentration was held constant for all measurements. To ensure that more than ∼95% of beads were driven by single dynein motors, the protein mixture was diluted before incubating with beads such that a maximum of 30% of beads exhibited activity when brought into contact with an axoneme (Belyy *et al*, 2016).

Optical trapping experiments were performed on a custom-built optical trap microscope set-up. DDB-bound beads were trapped with a 2 W 1,064-nm laser beam (Coherent) that was focused on the image plane using a 100x/1.49 N.A. oil-immersion objective (Nikon). Axonemes were located by brightfield imaging and moved to the center of the field-of-view with a locking XY stage (M-687, Physik Instrumente). Trapped beads were lowered to the axoneme surface with a piezo flexure objective scanner (P-721 PIFOC, Physik Instrumente). The position of the bead relative to the trap center was monitored by imaging the back-focal plane of a 1.4 N.A. oil-immersion condenser (Nikon) on a position-sensitive detector (First Sensor). A pair of perpendicular acousto-optical deflectors (AA Opto-Electronic) was used to control beam steering. To calibrate the detector response, a trapped bead was rapidly raster-scanned by the acousto-optical deflector and trap stiffness derived from the Lorentzian fit to the power spectrum of the trapped bead. The laser power was adjusted with a half-wave plate on a motorized rotary mount and maintained at a constant value of 80 mW, corresponding to a spring constant of 0.05 – 0.06 pN nm^-1^. Before data collection for each sample, the spring constant of a trapped bead was recorded.

In the case of DDB complexes formed with wild-type dynein, custom MATLAB software was used to extract stall forces and stall times from raw traces (following down-sampling from 5,000 Hz to 250 Hz). Stall events were defined as a stationary period of a bead at forces larger than 2.5 pN and durations longer than 100 ms, which were followed by snapping back of the bead to the trap center. Stall force was defined as the mean force during the last 20% of the stall event. The stall time was defined as the interval that the bead spent at a force of at least 80% of the stall force. All stall events were plotted and manually reviewed to confirm the accuracy of the computed values. Due to the distinct stall behavior of S3386C dynein, stall forces and stall times of this complex were calculated manually by reviewing each trace for stationary periods of a bead lasting longer than 150 ms at forces larger than 1 pN. The probability of dynein-driven beads sampling different forces under the trap was calculated by combining all the trajectories of wild-type or mutant dynein, plotting the normalized histogram of dynein at binned forces, and fitting the histogram to 3 Gaussians in MATLAB. The peak at near 0 pN represents the time the beads spend unbound from the microtubule. In Figure 9A, this peak was subtracted from the histograms to show the behavior of the beads when they are driven by dynein along the microtubule.

### Molecular dynamic simulations

The mouse cytoplasmic dynein-1 MTBD and partial stalk structure in complex with α/β-tubulin (PDB 6RZB; Lacey *et al*, 2019) was used to generate a model of the equivalent wild-type human cytoplasmic dynein-1 sequences bound to the tubulin heterodimer. Sequence alignment of the mouse and human MTBDs revealed no sequence divergence. However, two C-to-A mutations (at residues 3323 and 3387) had been introduced in the mouse MTBD used for cryo-EM and these were corrected in the human dynein model using the Mutator plugin of VMD (Humphrey *et al*, 1996). We used the same plugin to subsequently introduce the S3386C mutation into the model.

α-tubulin residues P37 to D47 are missing in PDB 6RZB, presumably because they are unstructured. A peptide was therefore constructed via the Molefacture plugin of VMD that has P37-D47 as an unstructured region and the flanking regions E23-M36 and S48-E55 as an α-helix and β-sheet, respectively. The constructed peptide was solvated in a water box and 150 mM KCl for MD simulations. The peptide system was minimized for 10,000 steps and followed by 2 ns of equilibration. After equilibration, the peptide was fitted into the remaining tubulin structure via targeted MD simulations (Schlitter *et al*, 1994) in which the flanking structured regions were slowly (10 ns) pushed into their crystal coordinates. Subsequently, the P37-D47 stretch was incorporated into the remaining tubulin structure using VMD’s merge extension.

MTBD-tubulin systems were aligned with the longitudinal axis of the α/β-tubulin dimer in the z direction. Each system was then solvated in a TIP3P water box with 35 Å cushions in each x-direction (70 Å water cushion in total) to provide enough space for stalk movements; 15 Å cushions were applied in all other directions. Systems were neutralized and ion concentrations set to 150 mM KCl. The size of the solvated system was ∼230,000 atoms. Simulations were performed at 310 K and 1 atm pressure with a time step of 2 fs. Langevin dynamics with a damping coefficient of 1 ps^-1^ were used to keep the temperature constant. The pressure was kept at 1 atm using the Langevin Nosé–Hoover method with a 100-fs oscillation period and a 50-fs damping time scale. Long-range electrostatic interactions were calculated with the particle-mesh Ewald method with a cut-off of 12 Å for van der Waals interactions. Prior to running MD simulations, 10,000 steps of minimization followed by 2 ns of equilibration were performed by keeping the proteins fixed. Subsequently, 10,000 steps of minimization were applied without any constraints on dynein and on tubulin, followed by 4 ns of equilibration with 1 kcal mol^-1^ Å^-2^ harmonic constraints on the C_α_. Starting from the equilibrated systems, 900-ns-long production runs were initiated with constraints on R2-H28, F53-E55, H61-D69, P72-R79, L92-S94, F103-N128, G131-N139, G144-Y161, S165-I171, V182-L195, V202-L217, Y224-S241, I252-L259, L269-A273, V288-F296, Y312-G321, P325-T337, F352-V353, R373-S381, R373-S381, and I384-H393 residues of α-tubulin and R2-H28, D41-T55, R58-D67, G71-A78, G82-F92, W101-G126, L130-S138, S145-L151, I163-V169, V180-S188, E198-M200, N204-T214, Y222-R241, G244-L246, L250-N256, F265-A271, V286-M299, Y310-G319, T323-N337, I349-C354, M363-S371, A373-Q391, R384-A393, and L395-G400 residues of β-tubulin to prevent structural deformation of the α/β-tubulin dimer due to the rest of the microtubule being absent. All MD simulations were performed in NAMD-2.14 (Phillips *et al*, 2020) using the CHARMM36 all-atom additive protein force field (Best *et al*, 2012).

To calculate MTBD angles in the plane perpendicular to the longitudinal microtubule axis, principal axes (PA) of microtubules were obtained using the VMD Orient tool. All PAs started from the origin located at the center of mass of tubulin and were referenced to the MTBD-tubulin structure. PA1 corresponds to the longitudinal axis of the protofilament. PA2 is the radial axis pointing towards the center of the A3295 and W3395 C_α_ atoms of dynein. PA3 is the tangential axis perpendicular to both PA1 and PA2. The MTBD vector, which indicates the combined MTBD and stalk base orientation with respect to the tubulin, was defined as the vector pointing from the center of dynein’s A3295 and W3395 C_α_ atoms toward the center of its A3288 Cα and Y3402 Cα atoms. To calculate the MTBD angle, the MTBD vector was projected onto the plane created by PA1 and PA2, and the angle between the projected vector and PA2 was evaluated.

We calculated the stalk angles relative to the microtubule longitudinal axis using the same procedure employed for the MTBD angles, but with a different selection of dynein atoms. Specifically, we superimposed the full-length monomeric dynein structure (PBD, 7Z8F; (Chaaban & Carter, 2022) with the MD conformations by using the Cα atoms of the stalk’s base, including residues A3295 to W3395 and A3288 Cα to Y3402, for alignment. We defined the stalk vector as the vector that originates from the center of dynein’s A3295 and W3395 Cα atoms, as sampled in the MD simulations, and extends towards the center of its R3191 Cα and S3501 Cα atoms within the superimposed structure.

The criteria for determining interactions were as follows. Salt bridge formations were detected using a maximum cut-off distance of 4 Å between the basic nitrogen and acidic oxygens (Barlow & Thornton, 1983). For hydrophobic interactions, a cutoff distance of 8 Å between side chain carbon atoms was used (Manavalan & Ponnuswamy, 1977; Stavrakoudis *et al*, 2009; Stock *et al*, 2015). To detect hydrogen bond formation, a maximum 3.5 Å distance between hydrogen bond donor and acceptor and a 30° angle between the hydrogen atom, the donor heavy atom, and the acceptor heavy atom was used (Durrant & McCammon, 2011).

### Statistics and data plotting

Statistical analysis and data plotting was performed with Prism 7.0b (GraphPad). Appropriate statistical tests were selected based on confirmed or assumed data distributions (Gaussian or non-Gaussian), variance (equal or unequal), sample size and number of comparisons (pairwise or multiple). Details of statistical tests applied for each dataset are provided in the relevant figure legends. Violin plots were used instead of bar charts when each group within a dataset had at least 30 values.

---

**Figure S1.**
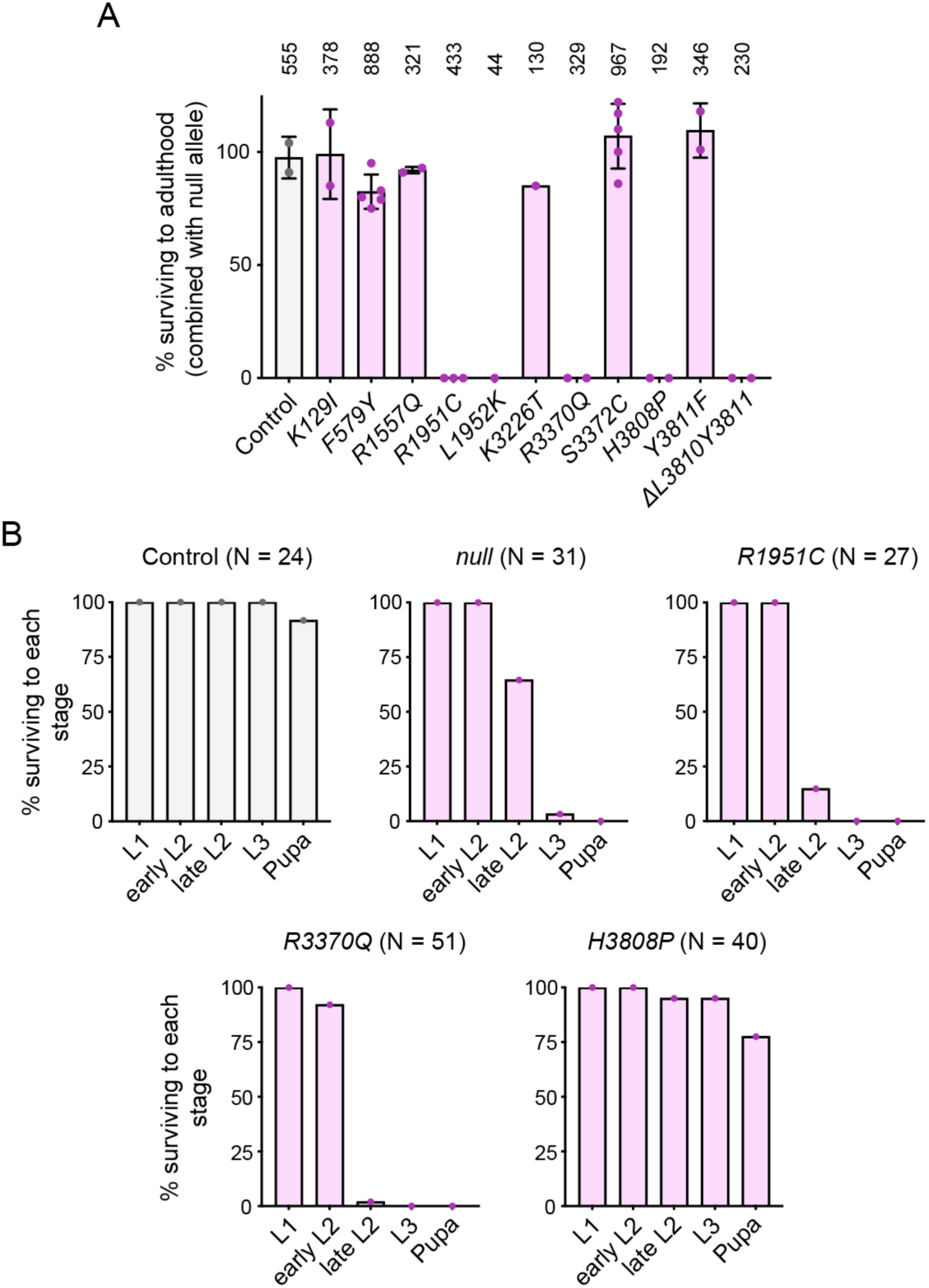
Lethality analysis of Dhc mutations. (A) Complementation tests showing percentage survival to adulthood of flies that are *trans-*heterozygous for the indicated *Dhc* mutation and a *Dhc^null^* allele. Columns display mean values from individual crosses; error bars are S.D.; circles are values for individual crosses. Data were normalized based on number of each genotype expected, if there were no lethality, given the total number of offspring. Numbers above each column show total number of offspring assessed for each genotype. (B) Percentage of L1 larvae of indicated genotypes reaching the specified stage (note that there was no overt lethality at the embryonic stage). N is number of animals analyzed for each genotype. In A and B, control animals were *trans*-heterozygous for a wild-type *Dhc* allele (recovered from the same CRISPR-Cas9 mutagenesis experiment that generated the *Dhc* mutant alleles) and the *Dhc^null^* allele.

**Figure S2.**
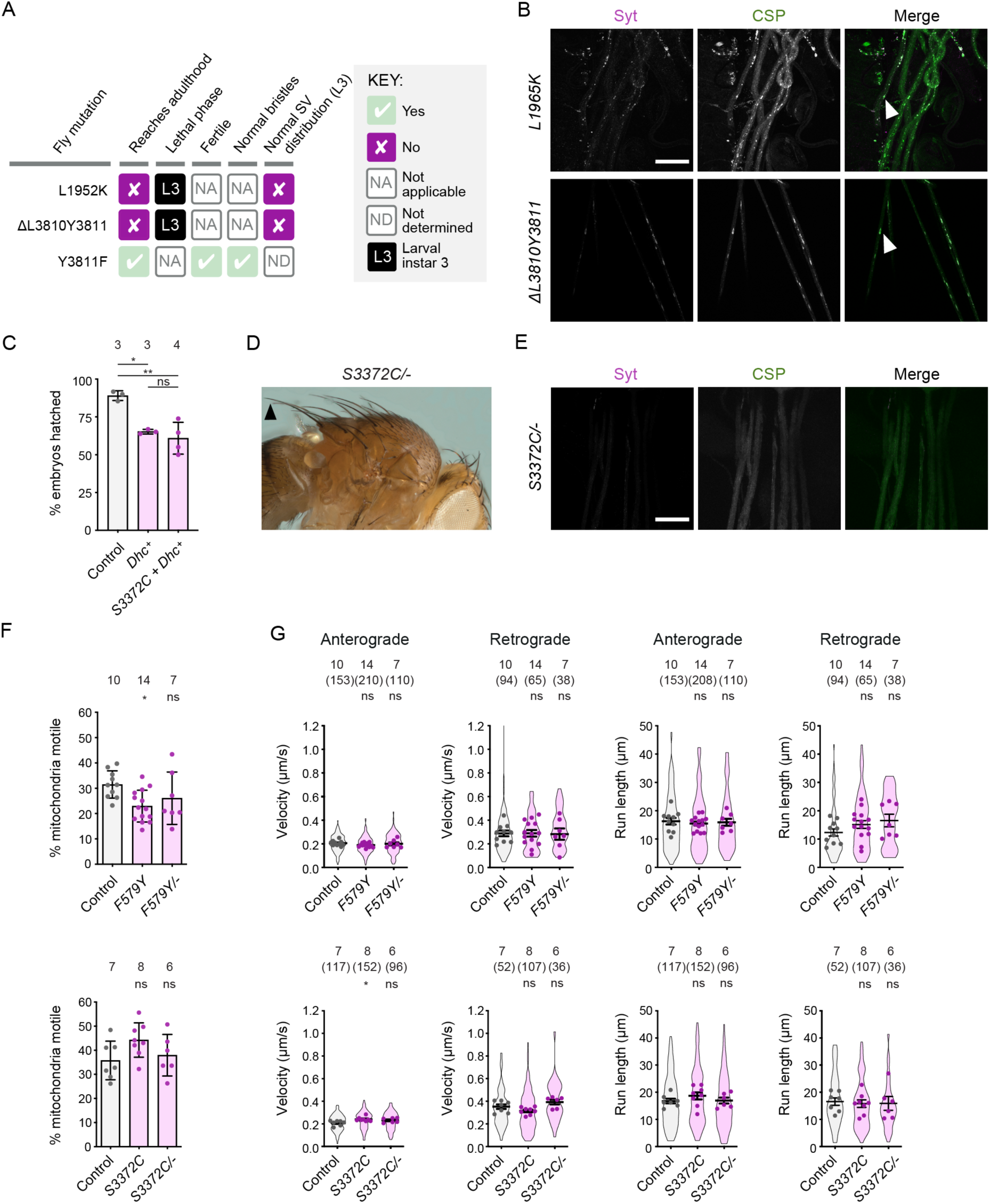
Supplementary data from phenotypic analysis of novel Dhc mutations. (A) Summary of *in vivo* effects of L1952K, ΔL3810Y3811 and Y3811F. SV, synaptic vesicle; L3, larval instar 3. (B) Confocal images of segmental nerves (taken proximal to the ventral ganglion; anterior to the top; Z-projection) from L3 larvae stained for the synaptic vesicle proteins Synaptotagmin (Syt) and Cysteine-string protein (CSP). Arrowheads show examples of synaptic vesicle accumulations in mutants. Images are representative of 3 – 6 larvae analyzed per genotype. See Figure 1E for images from control. (C) Quantification of hatching rate of eggs laid by mated females of indicated genotypes. Columns show mean values per egg collection; error bars represent S.D.; circles are values for individual egg collections. Number of collections per genotype (each from an independent cross; 414 – 1107 eggs per collection) is shown above bars. The control genotype was *yw*. *Dhc^+^* is a genomic rescue construct. Note that this construct reduces hatching rate in the wild-type background and that fertility defects of *S3372C* mothers are only suppressed by the transgene to this point. (D) Image (representative of >160 flies examined) showing normal bristles in *S3372C/-* adult flies. Arrowhead points to posterior scutellar macrochaetae. See Figure 1D for control image. (E) Confocal images of segmental nerves (taken proximal to the ventral ganglion; anterior to the top; Z-projection) from fixed L3 larvae stained for Syt and CSP, showing lack of abnormal synaptic vesicle accumulations. Images are representative of 3 larvae analyzed. See Figure 1E for images from control. (F) Quantification of percentage of mitochondria that exhibit transport in any direction in the adult wing nerve during the 3 minutes of data acquisition. Columns show mean values per movie; errors bars represent S.D.; circles are values for individual movies, each from a different wing. Number of wings analyzed is shown above bars. (G) Quantification of velocity and run length of transported mitochondria in the adult wing nerve. Violin plots show values for individual mitochondria and circles show mean values per wing. Horizontal lines shown mean ± S.D. of values for individual wings. Numbers without parentheses above bars are number of wings, with numbers of mitochondria given in parentheses. Evaluation of statistical significance (compared to control) in C, F and G was performed with a 1-way ANOVA with Dunnett’s multiple comparisons test (in G, the mean values per wing were compared): **, P<0.01; *, P<0.05; ns, not significant. Scale bars: B and E, 50 µm.

**Figure S3.**
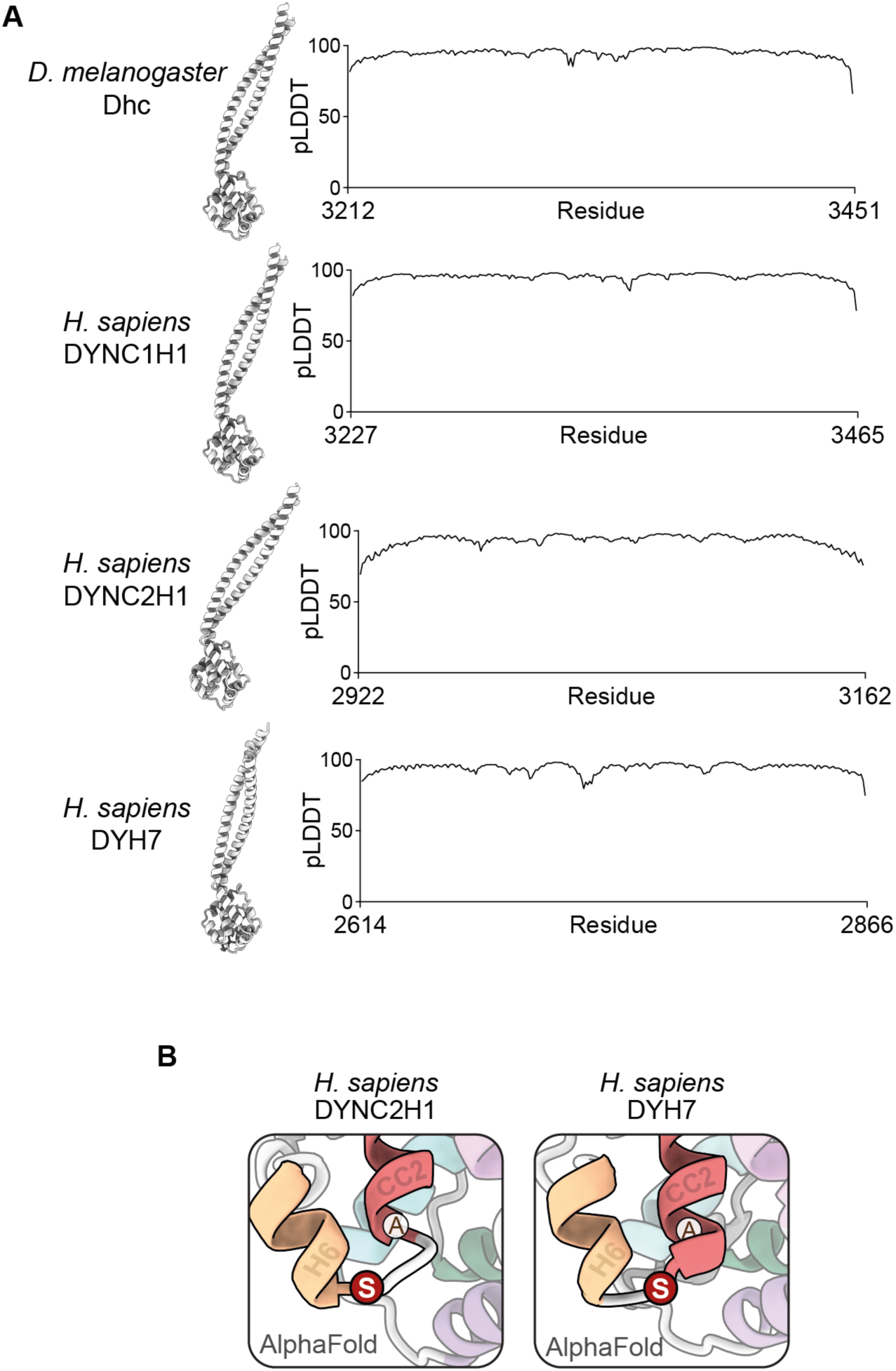
Supplementary data from structural analysis of S3372 in *Drosophila* Dhc and equivalent residues in other dynein family members. (A) Structural overview and pLDDT (predicted Local Distance Difference Test) plots (Jumper *et al*, 2021) of Alphafold2-generated structures of dynein MTBDs and stalks (showing that predictions are high confidence). (B) Zoom ins of regions contain serine residues equivalent to *Drosophila* S3372 in Alphafold2-generated structures of the MTBD and stalk of human DYNC2H1 (dynein-2 heavy chain) and DYH7 (inner arm axonemal dynein). Positions of residues equivalent to *Drosophila* S3372 are shown in red; alanines at residues equivalent to the cysteines at the base of CC2 in several other dyneins are also shown.

**Figure S4.**
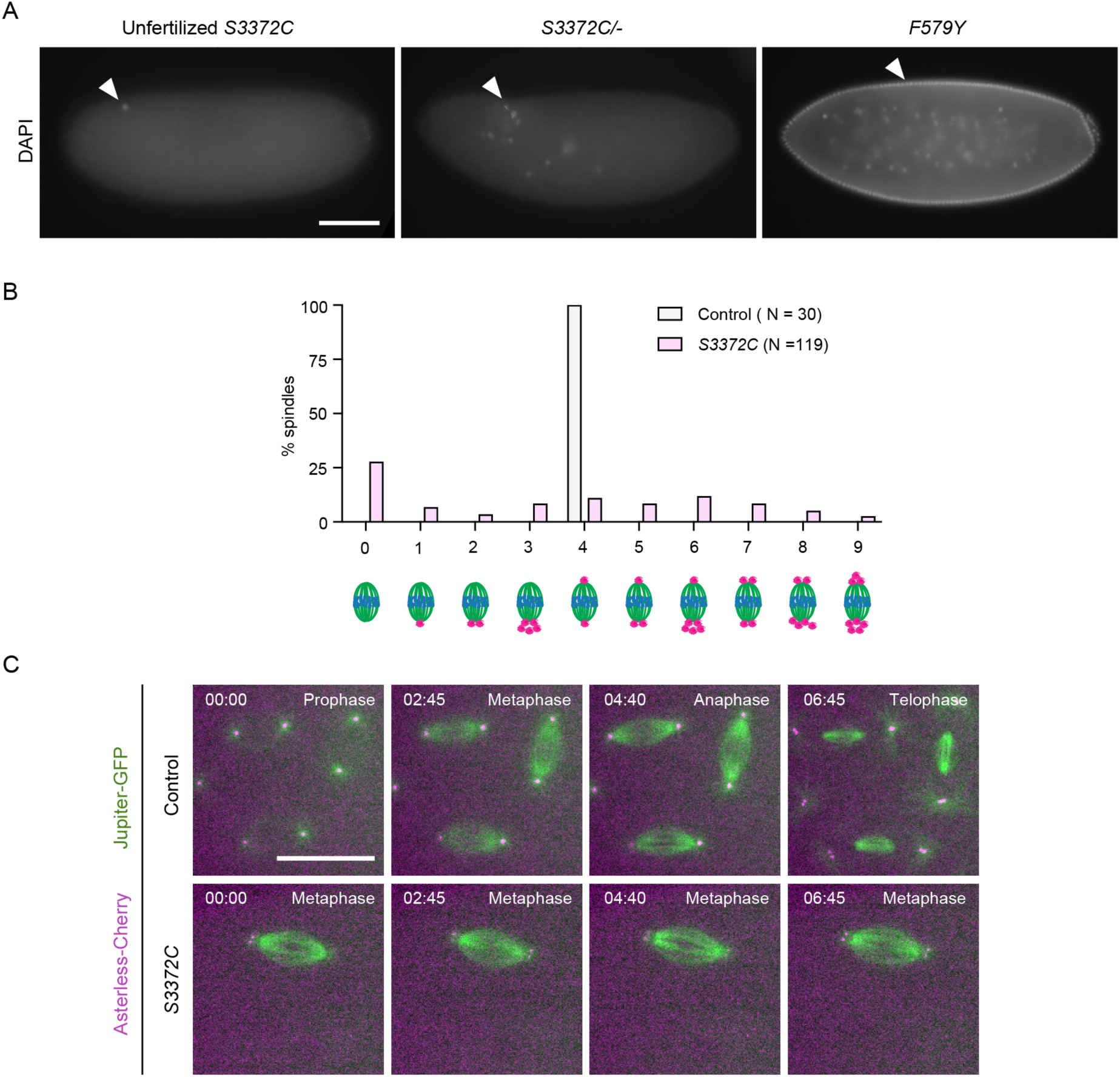
Supplementary data from coarse-grain analysis of S3372C mitotic phenotype. (A) Example wide-field images from a 2 to 4-h egg collection of fixed, DAPI-stained unfertilized eggs from virgin *S3372C* females, or embryos from mated *S3372C/-* or *F579Y* females (arrowheads show DNA staining). Images are representative of at least 150 embryos examined. (B) Categorization of mitotic spindle phenotypes in *S3372C* embryos based on centrosome number and arrangement. A range of mitotic stages were present in control embryos (*yw* strain), whereas >90% of mutant spindles were at metaphase; only those control and mutant embryos in metaphase were scored for this analysis. N is numbers of spindles scored (from 49 and 6 *S3372C* and control embryos, respectively). In both genotypes, no more than 5 randomly selected metaphase spindles were analyzed per embryo. (C) Example stills from time series (single focal plane) of control and *S3372C* embryos acquired during preblastoderm cycles. Jupiter-GFP and Asterless-Cherry label microtubules and centrosomes, respectively. Note abnormal presence of 2 centrosomes at each pole of the mutant spindle. In C, images were binned 2 x 2. Timestamps are min:s. Scale bars: A, 100 µm; B, 20 µm.

**Figure S5.**
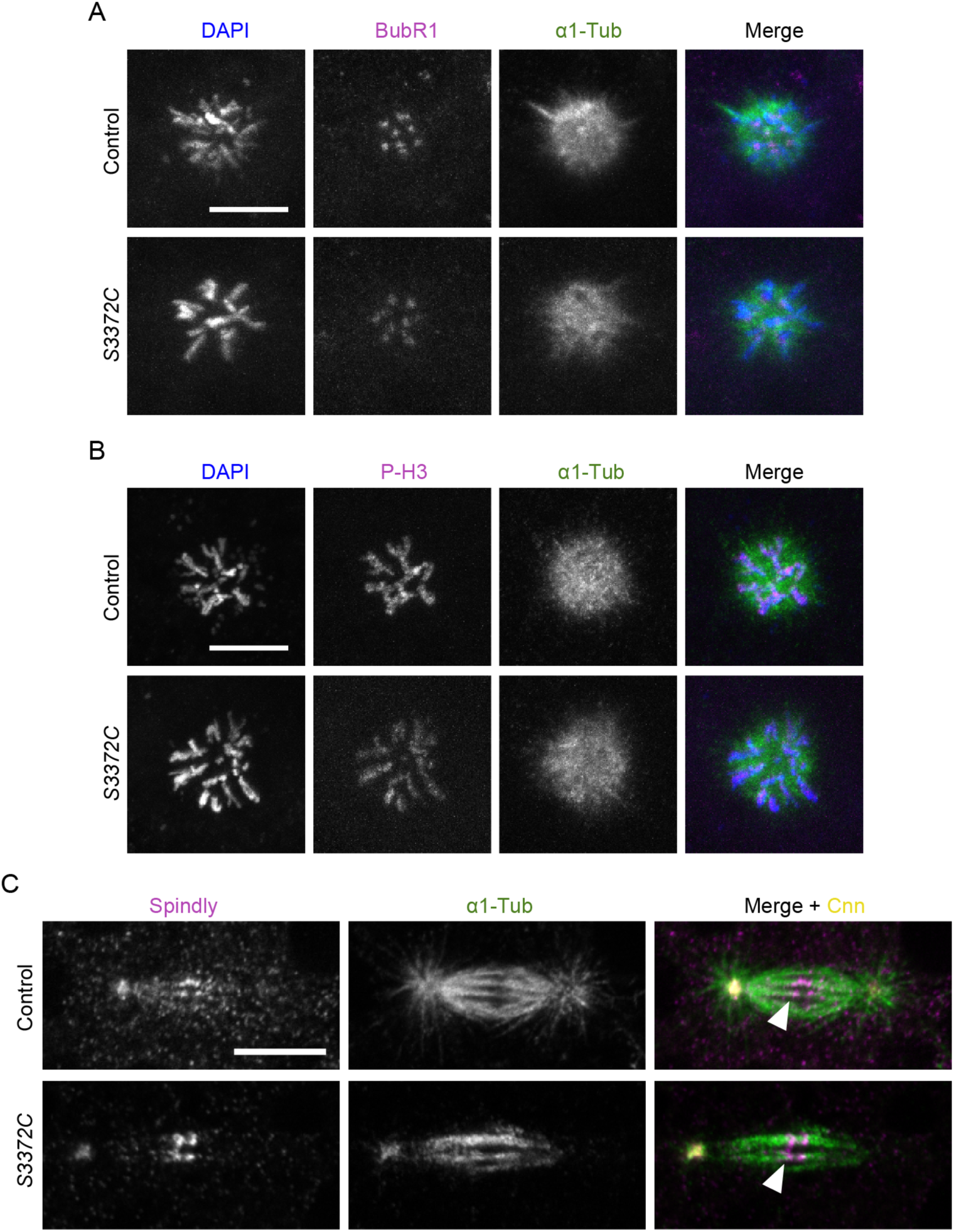
Supplementary data from fine-grained analysis of S3372C mitotic phenotype. (A, B) Example confocal images of polar bodies in fixed control and *S3372C* embryos stained with DAPI, as well as antibodies to (A) BubR1 and α1-Tubulin or (B) phospho-histone H3 (P-H3) and α1-Tubulin (Z-projections). (C) Example confocal images of mitotic spindles in control and *S3372C* embryos stained with antibodies to Spindly, α1-Tubulin and Centrosomin (Cnn), as well as DAPI. Arrowheads indicate example of close apposition of the ends of microtubule bundles and bright Spindly puncta at the kinetochore. Single focal planes were chosen to facilitate visualization of kinetochores, which resulted in only 1 centrosome being visible in each image. Timestamps are min:s. Scale bars in A – C, 10 µm. At least 50 embryos were examined per condition.

**Figure S6.**
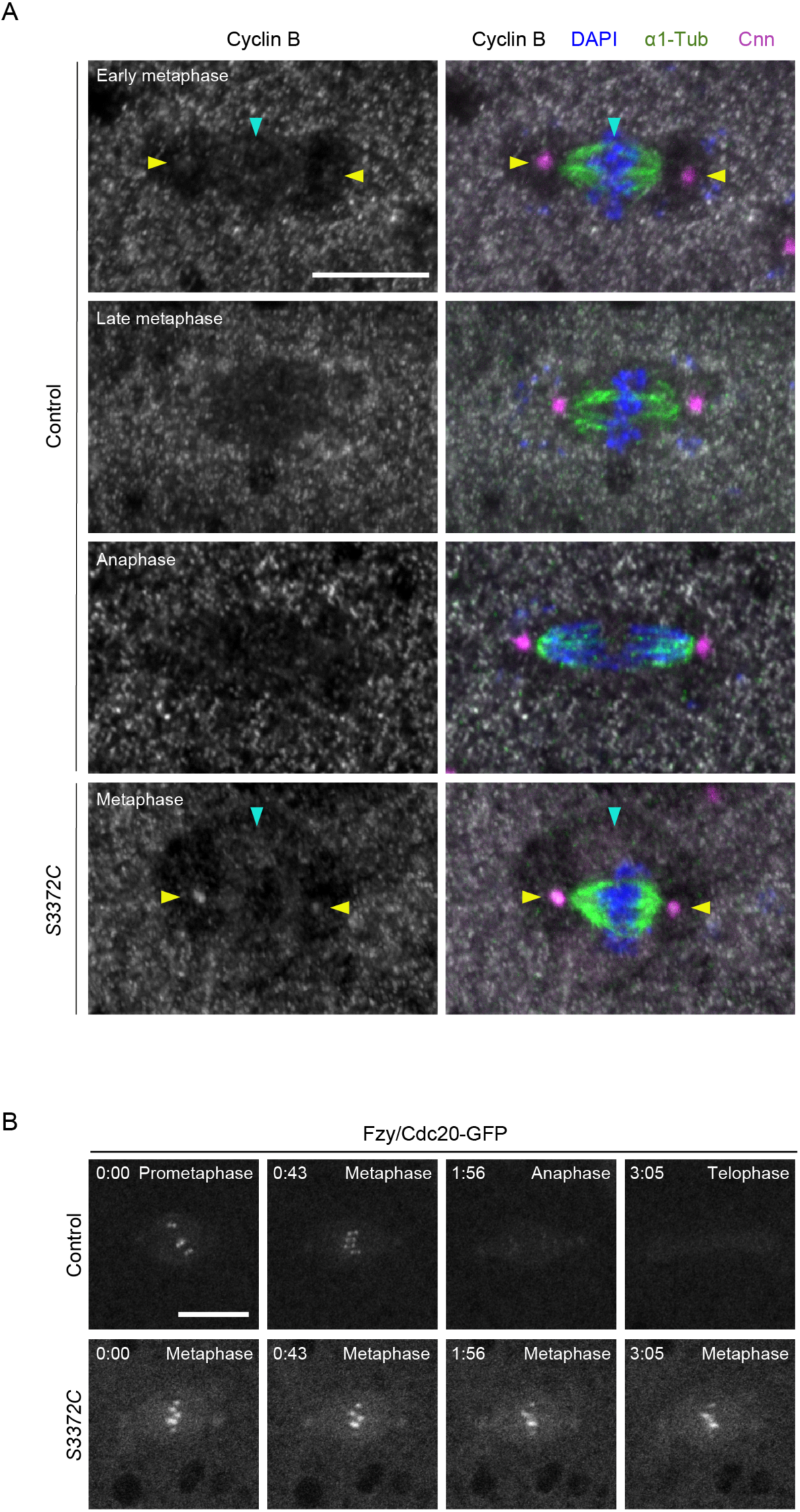
Analysis of Cyclin B and Fzy/Cdc20-GFP distribution on *S3372C* mutant spindles. (A) Example images of mitotic spindles during preblastoderm cycles in fixed control and *S3372C* embryos stained for the indicated antibodies, as well as with DAPI. Cyclin B shows weak accumulation in the vicinity of the spindle (blue arrowhead) and at centrosomes (yellow arrowheads) in early metaphase control embryos, which is lost by anaphase. In metaphase-arrested *S3372C* embryos, Cyclin B is detected in the vicinity of the spindle (blue arrowhead) and at centrosomes (yellow arrowheads). (B) Example stills of time series (single focal plane) of mitotic spindles acquired during preblastoderm cycles in live control and *S3372C* embryos expressing Fzy/Cdc20-GFP embryos. Fzy/Cdc20-GFP is localized to the metaphase plate in metaphase-arrested mutant spindles. Timestamps are min:s. Scale bars in A and B, 10 µm. At least 30 embryos were examined per condition.

**Figure S7.**
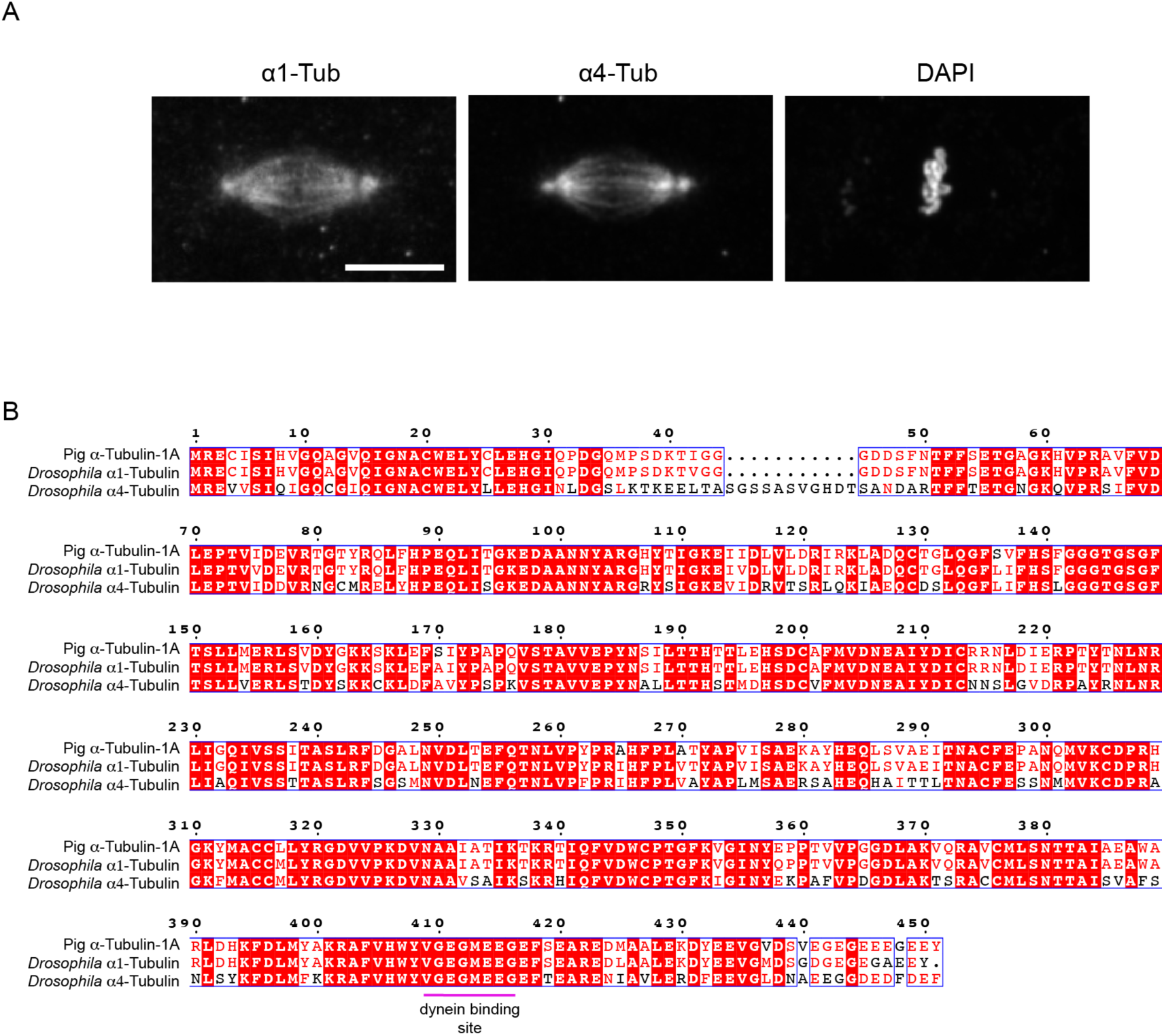
Assessing a potential α4-tubulin-specific effect of S3372C. (A) Example image of a metaphase spindle in wild-type embryo stained with antibodies to α1-tubulin and α4-tubulin, as well as DAPI. Scale bar: 10 µm. (B) Alignment of protein sequences of mammalian (pig) α-tubulin-1A, *Drosophila* α1-tubulin and *Drosophila* α4-tubulin. White letters on a red background indicate residues present in all sequences; red letters indicate residues present in ≥ 50% of sequences; blue boxes show regions with ≥ 50% conservation; magenta horizontal line, region contributing to dynein binding based on the mouse MTBD-microtubule structure (Lacey *et al*, 2019), which is identical in all 3 proteins. Uniprot accession numbers are: pig (*Sus scrofa*) α-tubulin-1A, P02550; *Drosophila melanogaster* α1-tubulin P06603; *Drosophila melanogaster* α4-tubulin P06606.

**Figure S8.**
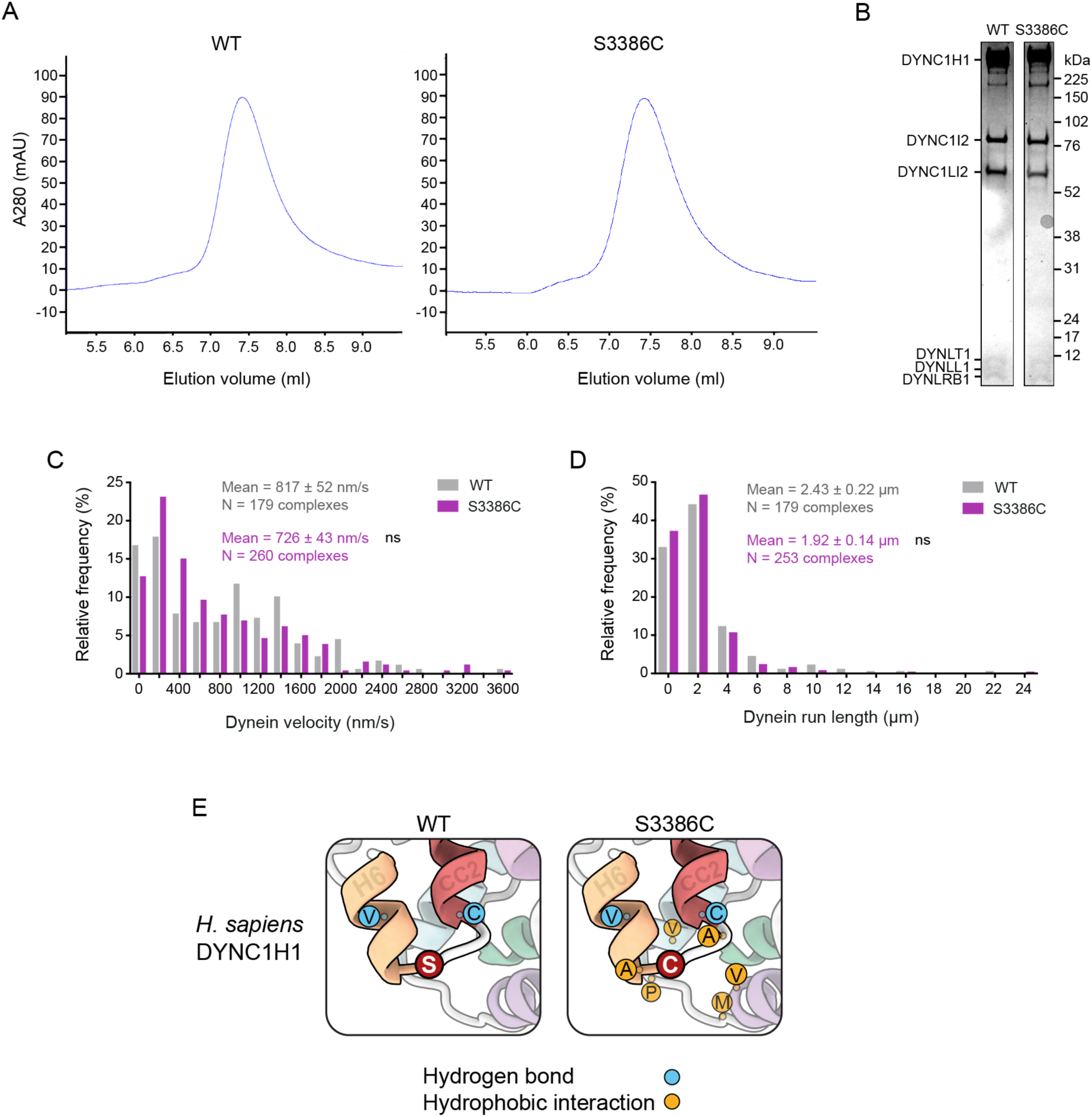
Supplementary data on *in vitro* and *in silico* analysis of S3386C human dynein. (A) Size-exclusion chromatography traces for wild-type (WT) and S3386C human dynein complexes, showing very similar profiles and lack of aggregation. (B) Cropped images from a Coomassie Blue-stained SDS-PAGE gel of pooled and concentrated fractions collected from the wild-type and S3386C mutant dynein peaks in A. (C, D) Velocity (C) and run length (D) frequency distributions for processive WT and S3386C mutant dynein complexes in the presence of dynactin and BICD2N in the assembly mix. Errors are S.E.M.. Evaluation of statistical significance was performed with a Mann-Whitney test. ns, not significant. (E) Zoom ins of regions of representative examples of MD-generated WT and S3386C mutant human dynein MTBD and partial stalk structures showing frequent hydrogen bonding interactions (blue circles) of S3386 and C3386 and new hydrophobic interactions of C3386 (gold circles); see Tables S1 – S3 for details of interacting residues.

## SUPPLEMENTARY MOVIE LEGENDS

**Supplementary movie 1. Live imaging of microtubules and chromatin in embryonic mitosis.** Composite of example movies (generated with spinning disk confocal microscopy) of a single focal plane of control and *S3372C* preblastoderm cycle embryos. Jupiter-GFP (green) and His2Av-mRFP (magenta) label microtubules and chromatin, respectively. Images were collected at 0.2 frames s^-1^ with a playback rate of 10 frames s^-1^. Timestamps are min:s. Scale bar, 10 µm. Related to Figure 5.

**Supplementary movie 2. Live imaging of microtubules and centrosomes in embryonic mitosis.** Composite of example movies (generated with spinning disk confocal microscopy) of a single focal plane of control and *S3372C* preblastoderm cycle embryos. Jupiter-GFP (green) and Asterless-Cherry (magenta) label microtubules and centrosomes, respectively. Images were collected at 0.2 frames s^-1^ with a playback rate of 15 frames s^-1^. Images were binned 2 x 2. Timestamps are min:s. Scale bar, 10 µm. Related to Figure S4.

**Supplementary movie 3. Live imaging of dynein and kinetochores in embryonic mitosis.** Composites of example movies (generated with spinning disk confocal microscopy) of a single focal plane of control and *S3372C* preblastoderm cycle embryos. GFP-Dlic (green in top merged movies) and Spc25-RFP (magenta in top merged movies) label dynein complexes and kinetochores, respectively. Two loops of the movie are shown; the second loop pauses to show transient accumulation of GFP-Dlic at kinetochores (arrows). Images were collected at 0.2 frames s^-1^ with a playback rate of 10 frames s^-1^. Timestamps are min:s. Scale bar, 10 µm. Related to Figure 6.

**Supplementary movie 4. Live imaging of GFP-Rod streaming in embryonic mitosis.** Composite of example movies (generated with spinning disk confocal microscopy) of a single focal plane of control and *S3372C* preblastoderm cycle embryos showing GFP-Rod dynamics. Streaming of faint GFP-Rod signals away from the kinetochores was analysis in kymographs. Images were collected at 0.5 frames s^-1^ with a playback rate of 10 frames s^-1^. Timestamps are min:s. Scale bar, 20 µm. Related to Figure 7.

## SUPPLEMENTARY TABLES

**Table S1.** Occurrence in simulations of hydrogen bond pairs involving S3386.

| **Donor (position)** | **Acceptor (position)** | **Occurrence (%) <sup>(1)</sup>** |
| --- | --- | --- |
| Ser3386 (loop) | Val3382 (helix 6) | 61.33 |
| Cys3389 (CC2) | Ser3386 (loop) | 41.24 |

(1) 3.6 $\mu$ s of all-atom MD simulations

**Table S2.** Occurrence in simulations of hydrogen bond pairs involving C3386.

| **Donor (position)** | **Acceptor (position)** | **Occurrence (%) <sup>(1)</sup>** |
| --- | --- | --- |
| Cys3386 (loop) | Val3382 (helix 6) | 69.20 |
| Cys3389 (CC2) | Cys3386 (loop) | 32.32 |

(1) 3.6 $\mu$ s of all-atom MD simulations

**Table S3.** Occurrence in simulations of hydrophobic interaction partners of C3386.

| Partner (position) | Occurrence (%) <sup>(1)</sup> |
| --- | --- |
| Val3307 (helix 1) | 99.94 |
| Met3310 (loop between helices 1 and 2) | 100 |
| Pro3314 (loop between helices 1 and 2) | 99.80 |
| Val3317 (helix 2) | 100 |
| Val3382 (helix 6) | 96.76 |
| Ala3385 (loop between helix 6 and CC2) | 100 |
| Ala3388 (CC2) | 100 |
| Cys3389 (CC2) | 100 |

(1) 3.6 $\mu$ s of all-atom MD simulations

**Table S4.**
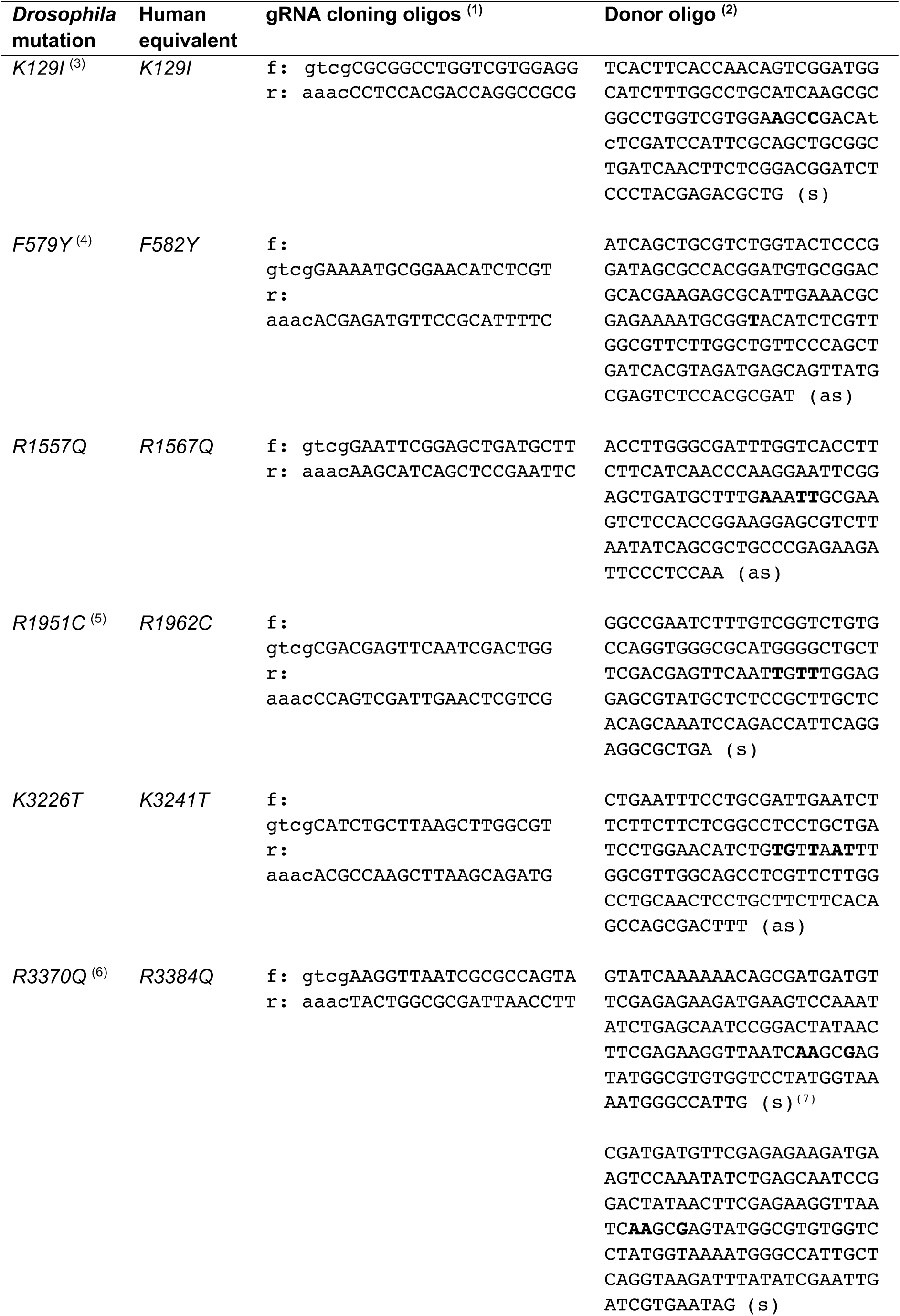
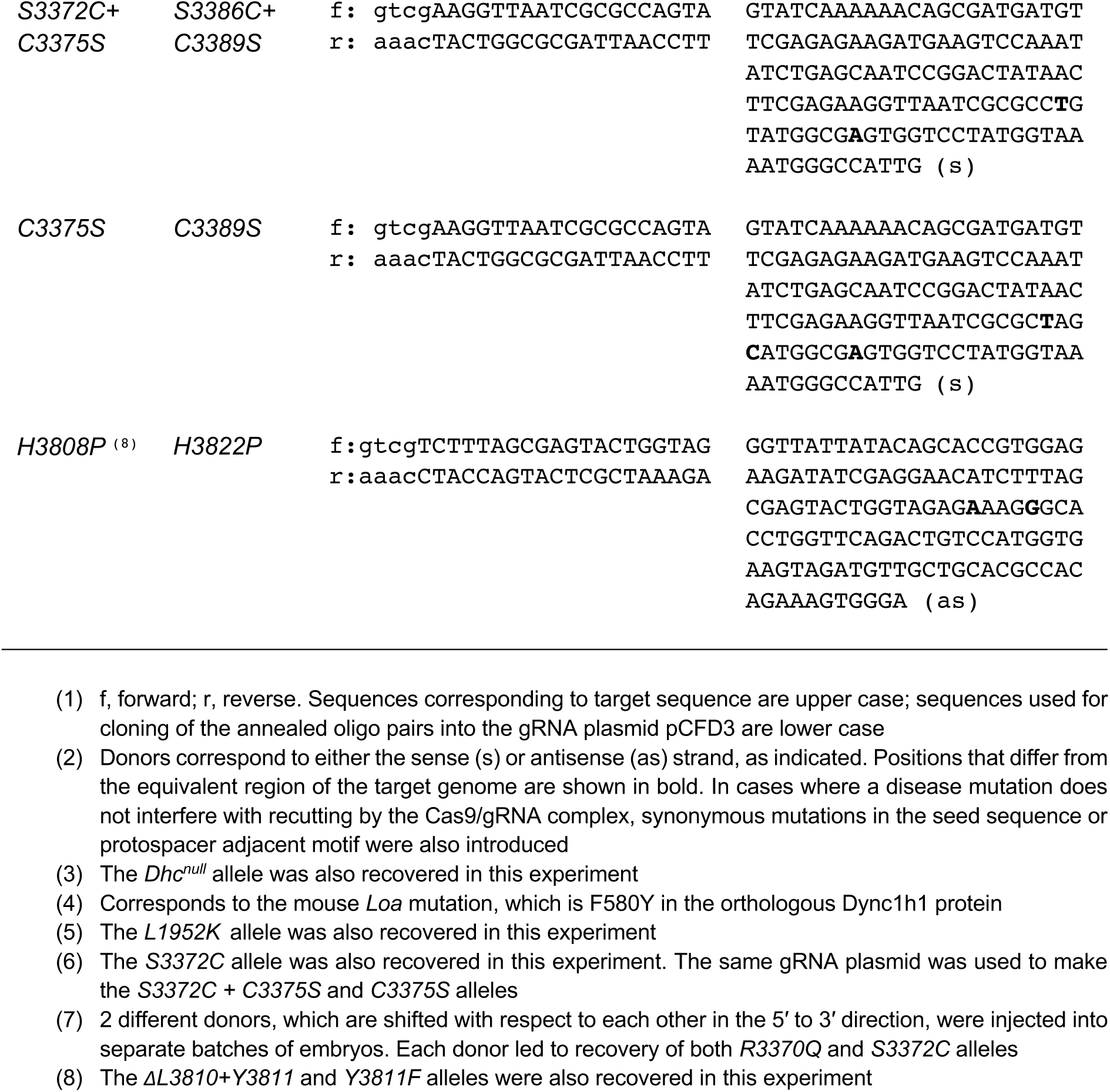
Oligonucleotides used to create Dhc mutations in *Drosophila*.

**Table S5.** Primary antibodies used in this study.

| Target | Host | Catalog number / Source <sup>(1)</sup> | Dilution for IF / IB <sup>(2)</sup> |
| --- | --- | --- | --- |
| α-Tubulin | Mouse | T6199 (clone DM1A) / Sigma Aldrich | 1:500 / NA |
| α-Tubulin | Rabbit | EP1332Y / Abcam | NA / 1:2000 |
| α4-Tubulin | Rabbit | NA / Fahmy *et al*, 2014 | 1:400 / NA |
| BubR1 | Rabbit | NA / Logarinho *et al*, 2004 | 1:1000 / NA |
| Centrosomin | Sheep | NA / Lucas & Raff, 2007 | 1:500 / NA |
| Cyclin B | Rabbit | NA / Whitfield *et al*, 1990 | 1:500 / NA |
| Cysteine string protein | Mouse | ab49 / Developmental Studies Hybridoma Bank (DSHB) | 1:250 / NA |
| Dynein heavy chain | Mouse | 2C11-2 / DSHB | NA / 1:1000 |
| Phospho-histone H3 | Rabbit | 9701 / Cell Signalling | 1:200 / NA |
| Spindly | Rabbit | NA / Griffis *et al*, 2007 | 1:1000 / NA |
| Synaptotagmin | Rabbit | NA / West *et al*, 2015 | 1:1000 / NA |

(1) NA, not applicable
(2) IF, immunofluorescence; IB, immunoblotting; NA, not applicable

**Table S6.**
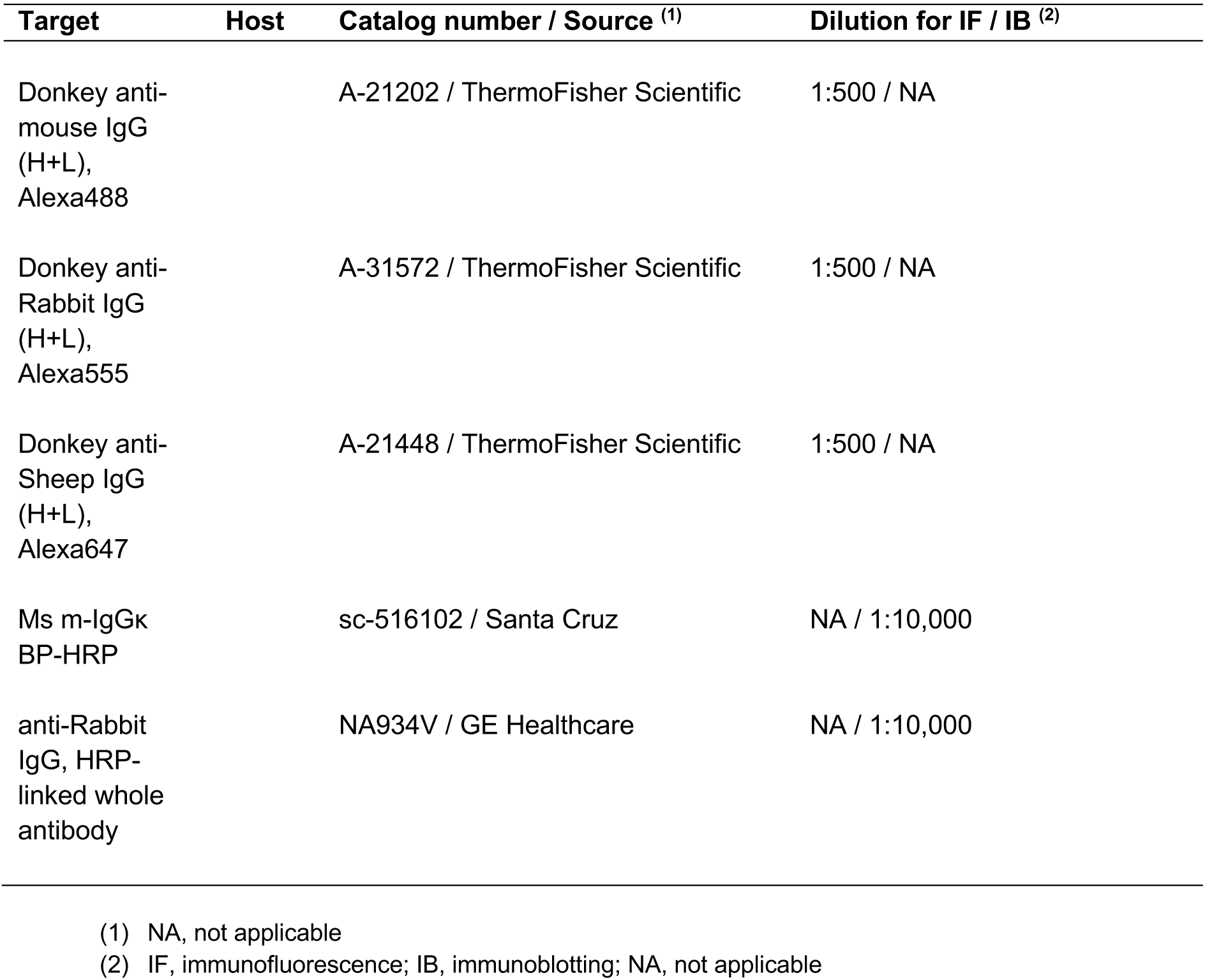
Secondary antibodies used in this study.

